# Learning the Language of Antibody Hypervariability

**DOI:** 10.1101/2023.04.26.538476

**Authors:** Rohit Singh, Chiho Im, Yu Qiu, Brian Mackness, Abhinav Gupta, Taylor Sorenson, Samuel Sledzieski, Lena Erlach, Maria Wendt, Yves Fomekong Nanfack, Bryan Bryson, Bonnie Berger

## Abstract

Protein language models (PLMs) based on machine learning have demonstrated impressive success in predicting protein structure and function. However, general-purpose (“foundational”) PLMs have limited performance in predicting antibodies due to the latter’s hypervariable regions, which do not conform to the evolutionary conservation principles that such models rely on. In this study, we propose a new transfer learning framework called AbMAP, which fine-tunes foundational models for antibody-sequence inputs by supervising on antibody structure and binding specificity examples. Our feature representations accurately predict an antibody’s 3D structure, mutational effects on antigen binding, and paratope identification. AbMAP’s scalability paves the way for large-scale analyses of human antibody repertoires. AbMAP representations of immune repertoires reveal a remarkable overlap across individuals, overcoming the limitations of sequence analyses. Our findings provide compelling evidence for the hypothesis that antibody repertoires of individuals tend to converge towards comparable structural and functional coverage. We validate AbMAP for antibody optimization, applying it to optimize a set of antibodies that bind to a SARS-CoV-2 peptide and obtaining 82% hit-rate and upto 22-fold increase in binding affinity. We anticipate AbMAP will accelerate the efficient design and modeling of antibodies and expedite the discovery of antibody-based therapeutics.

**Availability:**https://github.com/rs239/ablm

---

In modern therapeutics, antibodies have been some of the most promising drug candidates (*1*). This therapeutic success has been due to the remarkable structural diversity of antibodies, allowing them to recognize an extremely wide variety of potential targets. This diversity originates from their hypervariable regions which are critical to the functional specificity of antibodies. Unfortunately, deciphering how the sequence of an antibody’s hypervariable regions maps to its structure and function remains a gap in the field. Efficiently decoding this mapping is also crucial to a deeper understanding of human humoral immunity, as it will unlock characterization and comparison of immune repertoires across individuals and disease conditions.

Traditionally, experimental design of an antibody against a target of interest has been done by approaches like immunization or with directed evolution techniques like phage display selection (*2*). However, these methods have their limitations. The generation and screening process is slow and expensive. It also does not systematically explore the possible structural space, potentially leading to candidates with suboptimal binding characteristics. Furthermore, downstream considerations (e.g., developability or function-specific engineering) can not be easily accommodated (*3*). To address these challenges, there is a need for computational methods that can design a new antibody from scratch given a target (*4*) or which can more efficiently refine a small set of experimentally-determined candidates.

AlphaFold 2 (*5*) and related machine learning models (*6,7*) have demonstrated breakthroughs in the general task of protein structure prediction. However, they can struggle to predict antibody structures since the latter’s hypervariable regions (also known as the complementarity determining regions, CDRs) display evolutionarily novel structure patterns. One approach to tackle this issue has been to model the 3D structure of the entire antibody, or just its CDRs (*8,9*), but these have had limited accuracy. Moreover, they require many minutes per antibody (or CDR) structure, making it infeasible to perform large-scale computational exploration or analyze an individual’s antibody repertoire, which may contain millions of sequences.

In recent years, the advent of single-cell techniques for sequencing B-cell receptors has enabled large-scale data collection on immune repertoires (*10*). Leveraging these repertoires to inform individual and disease-level investigations of the humoral immune system requires analytic approaches that go beyond traditional clonal analysis to capture structure-function convergence. However, existing 3D structure-based approaches cannot scale to these large datasets, which can contain millions of sequences per individual.

As an alternative to explicit structure prediction, machine learning techniques used in natural language processing have been applied to generate high-dimensional protein representations (*11, 12, 13, 14, 15*). Protein language models (PLMs) capture structural features implicitly and also enable protein-property prediction. In the context of antibodies, one approach is to simply use PLMs trained on the corpus of all proteins (e.g., ESM (*11*)). We refer to these as “foundational” PLMs, the machine learning term for broad, general-purpose models (*16*). However, the CDRs of antibodies explicitly violate the distributional hypothesis underlying foundational PLMs: sequence variability in CDRs is *not* evolutionarily constrained. Indeed, the corresponding lack of high-quality multiple sequence alignments (MSAs) for antibodies is a key reason why AlphaFold 2 works less well on them than regular proteins (*5*).

To overcome the limitations of foundational PLMs, another set of approaches (e.g., AntiBERTa (*17*), IgLM (*18*)) has been proposed: these train the PLM just on antibody and B-cell receptor sequence repertoires. While these approaches better address the CDRs’ hypervariability, they have the disadvantage of not being trained on the diverse corpus of all protein sequences and are thus unable to access the substantial insights available from foundational PLMs (*12, 11, 14*). Moreover, existing approaches like AntiBERTa expend valuable explanatory power on modeling also the non-CDRs of the antibody, which are not very diverse and substantially less crucial to antibody binding-specificity. Lastly, neither set of approaches takes advantage of the 3D structures available for over 6,700 antibodies in the PDB.

Here, we argue that a more effective approach is to combine the strengths of the two approaches. We present a transfer learning approach that starts with a foundational PLM but adapts it for improved accuracy on the hypervariable regions by training on antibody-specific corpora. Such training lets us take advantage of available antibody structures as well as highthroughput single-cell assays of antibody binding specificity. Moreover, this approach can also easily be ported to new foundational PLMs as they are introduced (e.g., ESM-2 (*7*) instead of ESM-1b).

We introduce AbMAP (Antibody Mutagenesis-Augmented Processing), a scalable transferlearning framework that is applicable to any foundational PLM, and unlocks greater accuracy in the prediction of an antibody’s structure and its key biochemical properties. Our broad conceptual advance is to address the weakness of foundational PLMs on antibody hypervariable regions by a supervised learning approach that is trained on antibody structure and binding-specificity profiles. Specifically, we introduce three key advances: a) maximally leverage available data by focusing the learning task only on antibody hypervariable regions; b) a contrastive augmentation approach to refine the baseline PLM’s hypervariable region embeddings; and c) a multi-task supervised learning formulation that considers antibody protein structure as well as binding specificity to obtain the representation. Since the function of an antibody is determined primarily by its hypervariable region, we focus on modeling this region (and its immediate framework neighborhood) rather than the full sequence.

We explored the potential of AbMAP for speeding up antibody design. Applying it on Engelhart et al.’s dataset on antibodies binding to a 15-mer peptide from the HR2 region of SARS-CoV-2 spike protein (*19*), we sought to efficiently design a diverse, high-confidence set of positive and control predictions. Towards this, we designed a computational framework for in silico candidate generation that we believe will be broadly useful. Measured using surface plasmon resonance assays, AbMAP demonstrated 82% hit-rate in predicting the relative affinity of de novo candidates towards the peptide antigen.

Towards a deeper understanding of humoral immunity, AbMAP enables fast, structure-aware analysis of antibody repertoires across individuals. An advantage of AbMAP is that isotype switching in the human body (where a heavy chain’s constant region is replaced while preserving its hypervariable region) makes its CDR-focused representation well-suited for characterizing CDR-driven antigen specificity in a repertoire. Analyzing Briney et al.’s data on the antibody repertoires of multiple individuals (*10*), we find that AbMAP embeddings across individual repertoires are in substantially greater agreement than indicated by traditional sequence alignment-based metrics. This suggests that similar binding profiles are being activated in each individual despite sequence diversity. The embedding space also provides a principled framework to evaluate candidate antibody sequences for drug-like characteristics.

We also applied AbMAP representations to a number of downstream tasks: identifying structural templates, predicting the binding energy changes (ΔΔG) resulting from mutations, and identifying the antibody’s paratope. We formulate structure prediction with AbMAP as a template-search task where, for a query antibody, we search a template database of AbMAP antibody embeddings. Even without template refinement, our predictions compared well with many state-of-the-art structure prediction techniques, both general (e.g., AlphaFold 2 (*5*)) and antibody-specific (e.g., DeepAb (*8*)). AbMAP especially excels on the prediction of individual CDR structures, which is critical for accurate design. Finally, we also applied AbMAP to identify antibodies that can neutralize more than one SARS-CoV-2 variant; its predictions are substantially more accurate than those gleaned from ProtBert or ESM-1b directly.

## AbMAP: a language model of antibody hypervariable regions

For a given antibody sequence, we start from an embedding generated by a foundational PLM and refine it with AbMAP. There are three main steps in our refinement: identification of CDRs, augmenting the foundational PLM embeddings to focus on the CDRs, and an attention-based fine-tuning of the embedding to better capture antibody structure and function. We first apply ANARCI (*20*), a hidden Markov model approach, to demarcate the boundaries of CDRs. We use Chothia numbering (*21*) which leverages well-known canonical patterns of antibody structure (e.g., a disulfide bond that spans CDR-H1 and CDR-H2 (*22*), a Tryptophan residue located immediately after CDR-L1, (*23*) etc.) to identify CDRs with high confidence. We extend each ANARCI-reported segment by two residues on either side, allowing us to compensate for potential errors in ANARCI’s inference and also include the bounding framework residues. A similar design choice was made by Liberis et al. in Parapred (*24*). Next, we apply a procedure that we term “contrastive augmentation”: we perform *in silico* mutagenesis in the CDRs by randomly replacing a CDR residue with another amino acid and generate foundational PLM embeddings for each mutant. We then compute the augmented embedding as the difference between the embeddings of the original sequence and the average over mutants. Our augmentation aims to subtract away the sub-space of the embedding that does *not* correspond to the CDRs and, akin to masked language modeling, it highlights the contribution to embedding of a specific amino acid by contrasting it against potential replacements. We then optimize this augmented embedding with a Siamese neural network architecture with a single transformer layer that takes pairs of antibody sequences as inputs. Fig. 1b provides an overview of the AbMAP architecture. We seek a final representation for each antibody where Euclidean distances capture structural and functional information. Towards this, we adopted a multi-task learning formulation, requiring that the embeddings capture both structural similarity (from SABDab (*25*)) as well as functional similarity (antigen binding specificity scores from Setliff et al.’s profiling of B-cell receptors (*26*)). Additionally, the per-task fixed-length embeddings, which are *L*_2_-normalized, are regularized using a max-entropy formulation so that all dimensions of the embedding space are spanned by learned representations (**Methods**). The AbMAP approach can be applied to any foundational PLM. Here, we have applied it on the three foundational PLMs: Bepler & Berger (*12, 13*), ESM-1b (*11*), and ProtBert (*14*), producing AbMAP representations that we denote as AbMAP-*{*B, E, P*}* respectively.

**Figure 1:**
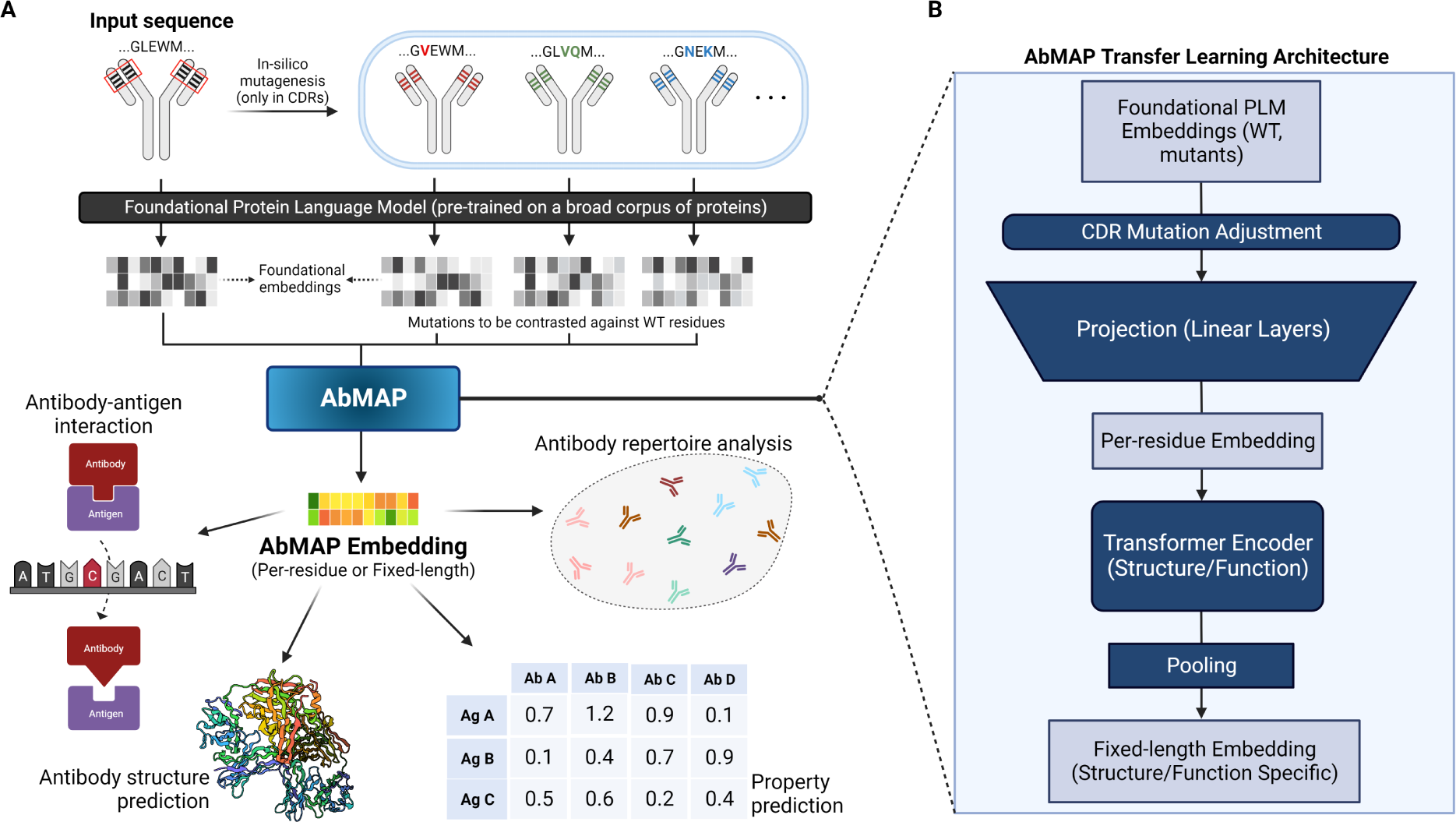
Overview of AbMAP embedding generation and its architecture. A) Given an input antibody sequence, our pipeline generates an embedding that can be applied for various downstream tasks including structure/property prediction as well as antibody repertoire analysis. B) AbMAP architecture comprises a projection module that applies contrastive augmentation and reduces the dimensionality of the input foundational PLM embedding to generate a variable length embedding, and a Transformer Encoder module that creates a *{*structure/function*}*-specific fixed-length embedding.

## AbMAP fine-tunes antibody embeddings effectively

We first assessed the effectiveness of our refinement and fine-tuning approach. After a random selection of 785 antibodies with available structures that were not in AbMAP’s training or validation set, we generated from them 10,000 random pairs and assessed how pairwise structural similarity correlated with representation similarity. We chose to evaluate structural similarity over the whole Fv, rather than just the CDR fragments, since we thought it to be a stricter test of the thesis that CDR-specific drivers are the primary determinants of overall structural variability across Fv structures. We evaluated representations from the baseline foundational PLM and each step of our refinement scheme: a) baseline (i.e., foundational) PLM representation for the whole protein, b) baseline PLM just for CDRs, c) with contrastive augmentation on CDRs and, d) also with the supervised transformer layer (residue-wise averaging for a-c). For each representation, we grouped the antibody pairs into 20 bins by cosine similarity and, in each bin, computed the distribution of TM-scores of structural similarity between the pairs. We assessed the embedding–structure relationship on consistency (measured as the Spearman rank correlation between average TM-score and cosine similarity across bins), as well as discriminative power (measured as the TM-score difference between the first and last bin). We show the results of AbMAP-B, (i.e. our method applied to the Bepler & Berger embedding) on heavy chain antibodies in Fig. 2. While the baseline foundational PLM is quite powerful on its own, it has somewhat limited consistency, especially on pairs with lower representation similarity. The CDR-specific embedding and contrastive augmentation improve the consistency (with Spearman correlation increasing from 0.94 to 0.98), while the final supervised layer is crucial to achieving strong discriminative power, with the TM-score separation between the the 2.5^th^ and 97.5^th^ percentile increasing from 0.04 to 0.37. Thus, applying a CDR-specific representation, augmenting it contrastively, and fine-tuning it in a supervised setting can accentuate antibody structural features more comprehensively and accurately than raw whole-protein PLM representations.

**Figure 2:**
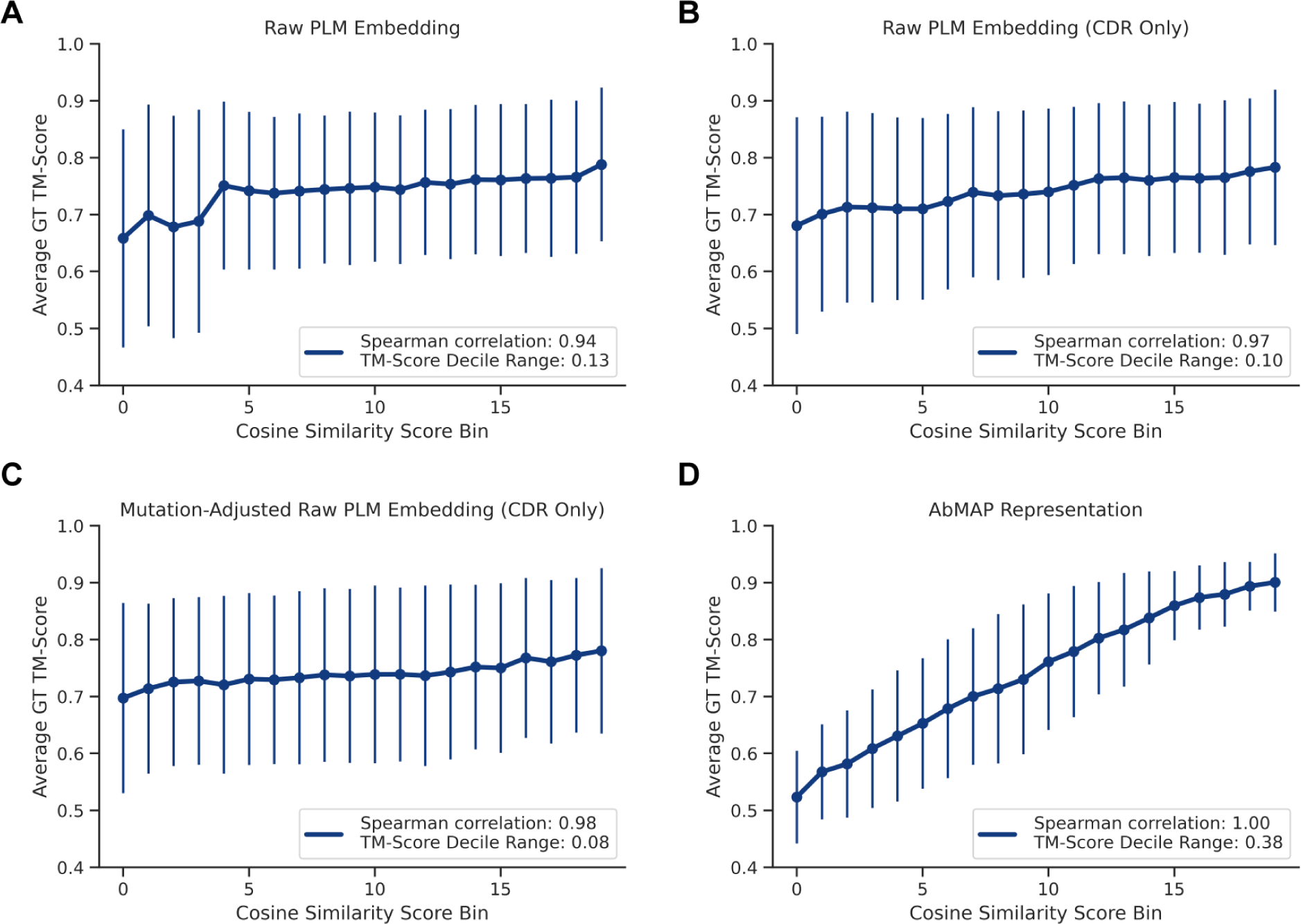
Average ground truth TM-scores antibody pairs in each of the 20 cosine similarity score bins. The embeddings used for cosine distance are A) raw PLM embeddings B) raw PLM embeddings on CDRs C) mutation-adjusted raw PLM embeddings on CDRs D) AbMAP fixed-length embeddings. Higher monotonicity in the plot implies that the model has successfully captured 3D structural information. Error bars indicate standard deviations.

## Antibody Structure Prediction

We approached structure prediction with AbMAP as a template-matching task: we searched through a database of antibody templates to find the example that we expect to be closest in structure to a query antibody. The template can later be refined (e.g., the Evoformer module (*5*)), a direction we hope to explore in future work (**Discussion**). For benchmarking purposes, the AbMAP template quality is itself an indication of the method’s ability to recapitulate structure. We constructed the template database from the set of SAbDab structures during AbMAP’s training; re-using these examples allowed us to evaluate on the remaining structures in SABDab. We first applied CD-HIT (*27,28*) to remove any template entries with greater than 70% sequence identity of test entries (our results are robust to this threshold as shown in Fig. 7). Using fixed-length AbMAP representations, we obtained the *k* (here, *k* = 10) templates closest to the query embedding by Euclidean distance (**Methods**). The medoid of these *k* representations

(in the Euclidean space) was reported as the matching template. Choosing the medoid instead of the very closest template offers some robustness to variability in embedding quality across queries and templates. As the underlying foundational PLMs improve, we expect this *k* could be lowered. We applied this process to generate templates from each foundational PLM as well as its AbMAP variants. For the former, the fixed-length embedding was obtained by taking the mean over the residues of the sequence’s PLM embedding. In addition to the foundational PLMs, we compared the performance of AbMAP against DeepAb, OmegaFold, and AlphaFold, some of the state-of-the-art deep learning-based methods for antibody structure prediction (*8, 6, 5*). To quantify the similarity between the predicted and ground truth structures, we computed the TM-scores (Template matching score) and RMSD (Root Mean Square Deviation) between the predicted and ground-truth Fv structures. While we consider both metrics, we believe TM-score is more appropriate since it is robust to variations in protein size, unlike RMSD. Since an antibody’s CDRs play crucial roles in its function, it is important that structure prediction models achieve high local accuracy on CDR structures. Accordingly, we also evaluated the methods on their predictions of specific CDRs. For individual CDR structure prediction, we prepared our embeddings separately, supervising with similarity scores for each CDR (H1-3, L1-3) instead of the whole structure.

Overall, as shown in Fig. 1, AbMAP (we show its -B variant here, with others reported in Table 3) is able to achieve high accuracy in structure prediction, despite no further refinement of the reported template. Compared to their respective foundational PLMs, each of the corresponding AbMAP variants performed substantially better. Overall, AbMAP-B performed better than other variants, possibly because the underlying foundation model is trained on both sequence and structure. Notably, AbMAP also improved over dedicated structure-prediction methods broadly. In particular, AlphaFold 2 performed substantially worse than others. This is consistent with reports of it under-performing on targets (like antibodies) where high-quality multiple sequence alignments are not available (*29*). AbMAP was also competitive with the language model-based OmegaFold, outperforming it on the TM-score metric and being roughly equivalent with it on the RMSD metric. On both metrics, the relative performance of AbMAP was especially strong on the crucial CDR-H3 region, suggesting that our generated embeddings may contain rich information about the CDRs that contribute the most to the antibody’s activity and specificity.

While the structures predicted by AbMAP can certainly be used for downstream tasks, we recommend directly using the PLM itself for downstream property-prediction tasks like paratope or ΔΔG prediction. Historically, explicit elucidation of the atomic coordinates of the antibody structure has been viewed as a prerequisite for such downstream tasks since they are informed by the structure’s physicochemical properties. However, we believe that the implicitness of representations encoded in a PLM like AbMAP allows a task-specific neural network greater power in marginalizing over unknowns and uncertainties in the structure (e.g., conformational flexibility); this implicit richness is lost when resorting to a single, fixed 3-D structure. Moreover, AbMAP offers the choice between a fixed-length embedding for property-prediction tasks and a per-residue variable-length embedding for tasks like in-silico mutagenesis; either embedding may be used as desired by the user.

## Variant Effect Prediction

A key application of computational antibody modeling is low-N antibody design and optimization: the task is to computationally extrapolate the effect of combinatorial mutations starting from a small training set of antibodies to a broad set of antibody candidates, using the results to guide the next round of assays. PLM-based *in silico* mutagenesis can play a key role in speeding up the design and development of antibody-based therapeutics. We assessed the generalization performance of AbMAP in estimating the binding efficacy of m396 mutants to SARS-CoV-2. The original wild-type variant of m396 targets the receptor binding domain (RBD) of the SARS-CoV-1 spike protein. During the COVID-19 pandemic, Desautels et al. sought to adapt this antibody to target the RBD of the SARS-CoV-2 spike protein. They generated 90,000 mutant in silico and estimated each mutant’s binding efficacy by computing its ΔΔG scores from five energy functions (FoldX Whole, FoldX Interface Only, Statium, Rosetta Flex, Rosetta Total Energy) (*30, 31, 32*). The effort required substantial high-performance computing resources from the Lawrence Livermore National Laboratory, and needed over 200,000 hours of CPU time (*33*). We sought to predict the ΔΔG scores for these set of mutants after training on as little as 0.5% of the examples. We evaluated two prediction architectures: i) using AbMAP’s variable-length embedding as input to a transformer layer followed by a two-layer feed-forward network (averaged over residues), and ii) using AbMAP’s fixed-length embedding as input to a ridge regression. The target variable in both cases was Desautels et al.’s binding efficacy score for the mutant. The first architecture offers greater explanatory capacity while the second ar-chitecture can be applied even with very limited training data. We trained these architectures using embeddings from AbMAP-B/E/P as well as their corresponding foundational PLMs. We also used a simple baseline embedding that uses a one-hot encoding of the amino acids. The intuition behind this baseline is that, given sufficiently many examples, the one-hot encoding can be leveraged effectively by the downstream predictor but its usefulness would diminish when fewer training examples are available. We trained the different models for 100 epochs each, and varied the train/test split ratio to examine how the performance degrades as fewer training examples are provided.

In our evaluations, we assessed both the overall accuracy of AbMAP-based predictions as well as its ability to recapitulate the top ground-truth hits. For the overall analysis, we computed the Spearman rank correlation between predicted and ground-truth scores, averaging these correlations over the five energy-function categories. As shown in Fig. 3a, with just 20% of the examples (i.e., train/test split of 0.2), AbMAP-E and AbMAP-P’s representations both achieve Spearman rank correlation of 0.94, indicating that AbMAP can effectively generalize from a limited training set, thus reducing experimental and computational expenses. In contrast, the raw PLMs perform substantially worse. Indeed, for the largest training set size (0.2 split), the lower-dimensional one-hot encoding performs better than the foundational PLMs. The performance comparison for AbMAP-B and the foundational Bepler & Berger PLM are shown in Fig. 9. Furthermore, the performance of AbMAP embeddings is more robust to smaller training set sizes than the baseline PLMs. The relative outperformance of AbMAP becomes more substantial as the number of training examples decreases. With just 0.5% of the examples (i.e., train/test split of 0.005), AbMAP is able to achieve high accuracy (Spearman rank correlations of 0.71 and 0.63 with the AbMAP-E and -P models, respectively). Notably, the ridge regression-based formulation starts to outperform the more complex model as the number of training examples decrease, as would be expected. High accuracy in such few-shot settings is crucial since they enable a small set of experimentally-assayed binding specificity/strength measurements to be extrapolated more broadly.

**Figure 3:**
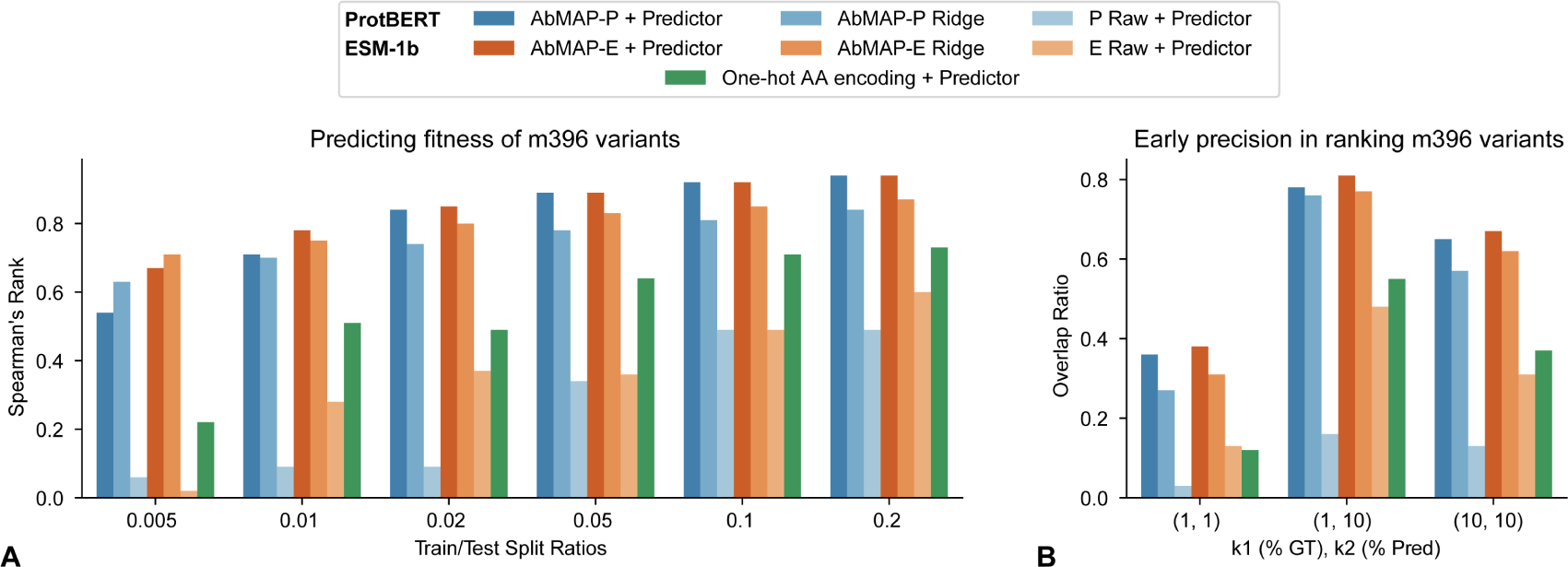
Comparative evaluation of protein representations for property prediction in a low-N context. For mutations of the wild-type SARS-CoV-1 antibody m396, methods were evaluated on their ability to extrapolate ΔΔG from limited training data, as well as the precision of their top predictions. AbMAP-*{*P,E*}* refer to AbMAP variants trained with **P**rotBERT and **E**SM models; the baseline models are “*{*P,E*}* Raw”. Each of these representations was used with a multi-layer perceptron (“Predictor”) to predict ΔΔG. To explore if AbMAP variants were informative enough for even simpler prediction heads, we also evaluated ridge regression-based prediction. **A**) Average Spearman correlation between ground-truth (GT) and predicted ΔΔG scores across various train/test splits. **B**) For a low-data setting (train: 0.02, test: 0.98), the fractional overlap between top ground-truth (*k*_1_) and predicted (*k*_2_) hits.

We also examined the early precision of our model, i.e., its ability to correctly prioritize mutants with high ground-truth scores. We checked how many sequences in the top *k*_1_% (*k*_1_ = 1, 10) of ground-truth scores overlapped with those in the top *k*_2_% (*k*_2_ = 1, 10) of the model’s predicted scores. Even at high cutoffs, when comparing the top 1% of predicted and ground truth scores by magnitude, there is 38% and 36% overlap when using AbMAP-E and AbMAP-P embeddings, respectively (Fig. 3b). Altogether, AbMAP-P/E are able to robustly predict mutational performance, both broadly and for the top hits. Crucially, they are able to operate much more effectively with limited training data, compared to the foundational PLM embeddings or one-hot encodings.

## Paratope Prediction

An important antibody sub-structure is the paratope – the region that recognizes and binds to an antigen. In particular, each residue on the antibody backbone can be assigned a binary label indicating if it belongs to the antibody’s paratope. We applied AbMAP-B to the paratope prediction task, comparing it against a dedicated machine learning method, Parapred (*24*), as well the ProtBert foundational PLM. Acquiring all SABDab heavy chain entries with at least one CDR in contact with an antigen, we labeled a residue as part of the paratope if it was within 5A^°^ of the antigen. To avoid data snooping, we re-used AbMAP’s training set for this task-specific training (1195 entries), and evaluated on the remaining test set of entries (312 entries). For paratope prediction, we specified a simple architecture that uses the per-residue, variable-length representation of AbMAP: a single transformer layer followed by two linear layers. Predictions from ProtBert were made using the same architecture. Notably, the ProtBert model has more parameters because the ProtBert embedding dimension (1024) is four times as large as AbMAP’s (256). The Parapred model takes individual CDR fragment strings as input, and we applied it separately on the three heavy chain CDRs. We calculated per-residue performance and report the overall statistics in Table 2. AbMAP-B achieves the highest overall accuracy for per-residue paratope prediction. The performance of Parapred reported here may be a slight overestimate since we use the trained model made available by Leem et al.’s PyTorch implementation (*34*); their training may have utilized some of the examples that we use here in the test set. While ProtBert has similar accuracy to AbMAP, it uses many more model parameters due to the larger embedding dimensionality.

**Table 1:** Comparison of AbMAP-B and other models for antibody structure prediction using both RMSD (C-alpha) and TM-Score as metrics. For AbMAP-B, ProtBert, and ESM-1b, the predicted template structure was selected from a set of antibodies whose sequence identity is below 0.7. Structure prediction was conducted on both individual CDR fragments and the whole Fv chain.

| Model | Metric | Chain | CDR 1 | CDR 2 | CDR 3 | Whole Fv |
| --- | --- | --- | --- | --- | --- | --- |
| AbMAP-B | TM-Score ↑ | H | **0.62** ± 0.007 | **0.67** ± 0.006 | **0.54** ± 0.007 | **0.86** ± 0.006 |
| ProtBert | | | 0.33 ± 0.003 | 0.31 ± 0.003 | 0.28 ± 0.003 | 0.69 ± 0.005 |
| ESM-1b | | | 0.49 ± 0.007 | 0.51 ± 0.006 | 0.44 ± 0.007 | 0.79 ± 0.004 |
| DeepAb | | | 0.51 ± 0.008 | 0.55 ± 0.008 | 0.24 ± 0.004 | 0.54 ± 0.005 |
| OmegaFold | | | 0.61 ± 0.007 | **0.67** ± 0.006 | 0.34 ± 0.005 | 0.79 ± 0.005 |
| AlphaFold2 | | | 0.28 ± 0.003 | 0.30 ± 0.003 | 0.30 ± 0.004 | 0.65 ± 0.004 |
| AbMAP-B | TM-Score ↑ | L | 0.62 ± 0.007 | **0.76** ± 0.007 | **0.65** ± 0.007 | **0.89** ± 0.004 |
| ProtBert | | | 0.34 ± 0.003 | 0.60 ± 0.007 | 0.52 ± 0.010 | 0.80 ± 0.005 |
| ESM-1b | | | 0.41 ± 0.005 | 0.61 ± 0.007 | 0.39 ± 0.005 | 0.82 ± 0.004 |
| DeepAb | | | 0.40 ± 0.006 | 0.66 ± 0.009 | 0.38 ± 0.006 | 0.52 ± 0.005 |
| OmegaFold | | | **0.63** ± 0.006 | 0.69 ± 0.006 | 0.58 ± 0.006 | 0.83 ± 0.005 |
| AlphaFold2 | | | 0.27 ± 0.003 | 0.36 ± 0.007 | 0.31 ± 0.003 | 0.58 ± 0.004 |
| AbMAP-B | RMSD ↓ | H | 0.43 ± 0.016 | 0.38 ± 0.013 | **0.43** ± 0.025 | 2.11 ± 0.082 |
| ProtBert | | | 1.40 ± 0.030 | 1.21 ± 0.028 | 0.64 ± 0.031 | 3.07 ± 0.078 |
| ESM-1b | | | 0.54 ± 0.017 | 0.76 ± 0.022 | 0.50 ± 0.023 | 2.69 ± 0.063 |
| DeepAb | | | 0.56 ± 0.016 | 0.55 ± 0.015 | 0.81 ± 0.031 | **0.72** ± 0.028 |
| OmegaFold | | | **0.35** ± 0.013 | **0.37** ± 0.013 | 0.75 ± 0.035 | 2.39 ± 0.067 |
| AlphaFold2 | | | 0.70 ± 0.032 | **0.37** ± 0.030 | 1.24 ± 0.063 | 4.40 ± 0.077 |
| AbMAP-B | RMSD ↓ | L | 0.41 ± 0.015 | 0.18 ± 0.006 | **0.44** ± 0.022 | 1.42 ± 0.044 |
| ProtBert | | | 1.39 ± 0.041 | **0.15** ± 0.008 | 0.64 ± 0.044 | **0.76** ± 0.012 |
| ESM-1b | | | 0.68 ± 0.022 | 0.22 ± 0.008 | 0.65 ± 0.028 | 2.38 ± 0.062 |
| DeepAb | | | 0.72 ± 0.021 | 0.32 ± 0.011 | 0.84 ± 0.030 | 1.16 ± 0.084 |
| OmegaFold | | | **0.40** ± 0.016 | 0.17 ± 0.007 | 0.46 ± 0.017 | 2.31 ± 0.076 |
| AlphaFold2 | | | 1.13 ± 0.046 | 0.24 ± 0.024 | 0.64 ± 0.033 | 4.79 ± 0.084 |

**Table 2:** Comparison of our model and ProtBert’s representations for antibody paratope prediction by training a separate predictor using the representations as input. The performance of our model’s representations was also compared against Parapred.

| Embedding | Dimension | Predictor | Accuracy | AUPRC | AUROC |
| --- | --- | --- | --- | --- | --- |
| AbMAP-B | 256 | 1-layer transformer | **0.798** | **0.653** | 0.830 |
| ProtBert | 1024 | 1-layer transformer | 0.797 | 0.646 | **0.842** |
| One-hot + Physicochemical | 28 | Parapred | 0.621 | 0.639 | 0.838 |

## AbMAP Enables Iterative Antibody Optimization

Antibody design presents a significant drug-design challenge, requiring precise optimization of affinity and specificity. AbMAP is designed to fit into existing workflows of iterative optimization, where candidates are progressively refined in each iteration. We explored how effectively AbMAP could improve the selection and refinement of antibody candidates, thereby contributing to a cheaper and more efficient design process. As a proof of concept, we optimized antibodies targeting a peptide in the HR2 domain of the SARS-CoV-2 spike protein, with data from Engelhart et al.’s study as a starting point (*19*). Comprising 104,972 antibodies from four libraries the study used yeast phage display technology to estimate the binding strength of each antibody, measured as the dissociation constant (kD). Each library was based on a wildtype antibody sequence and its set of mutants included all 1-point, many 2-point, and some 3-point mutations over the wild-type. We aimed to assess if we could expand the candidate set to explore additional 3-point mutations, as well as 4-point mutations. A secondary goal of our analysis was to establish a computational workflow for using embedding-based approaches like AbMAP for iterative optimization. Consequently, we sought to predict and validate not only strongly-binding mutants but also weakly-binding ones, both as a negative control and as valuable data for later rounds of optimization.

We employed AbMAP embeddings in classification and regression models trained to predict binding affinities from Engelhart et al.’s data. Our predictive framework is designed to address three key considerations frequently encountered in computationally-guided antibody optimization studies. First, the training data can come from multiple libraries, with substantial variations in binding efficacy across these libraries. In the original study, antibodies originated from four libraries (AAYL49, AAYL50, AAYL51, and AAYL52) each based on mutations of either the heavy or light chain of a starting wild-type antibody. Of these, the average binding strength, measured as log(kD), was the strongest for AAYL50. Hence, simply combining these libraries and predicting log(kD) as a regression target might overemphasize the contribution of the strongest library, failing to identify beneficial mutations in other libraries. Second, the typical design goal is to minimize kD, implying that predictive accuracy is more crucial at the lower end of kD predictions. Solely framing the task as a regression against log(kD) might inaccurately equate changes in the millimolar range with those in the nanomolar range. Third, diversity and robustness among candidates are highly desired. A diversity of candidates, covering distinct regions of the embedding space, provides a broader set of candidates for subsequent optimization rounds. For robustness, we focused on candidates where minor changes in embeddings would not significantly affect their binding characteristics.

We created an ensemble of predictive models. Using thresholds, we generated both positive (low kD) and negative (high kD) datasets. Two such models were developed, with thresholds of 10 and 100 nM respectively, and were trained using logistic regression for each library separately. We then computed ΔΔG scores for each mutant by adjusting against the library’s wild-type (i.e., ΔΔG = log(kD_m_*_utant_*) - log(kD_W_*_T_*)), reducing variability across libraries. After pooling all libraries, we trained a regression model. For each library, we generated 500,000 3-point and 4-point mutations. Candidates were scored by the three models, and those ranking in the top 5% for each model were shortlisted, yielding 700-1100 candidates per library. These were clustered using k-means clustering (k=20), with cluster centroids nominated as positive candidates. An analogous procedure was used for negative candidates, focusing on low-scoring ones.

We next sought to experimentally validate the predictions of AbMAP. We recombinantly produced 32 antibodies across the four libraries from the Engelhart et al. study (AAYL49, AAYL50, AAYL51, AAYL52). We synthesized the original wild type antibody for each library and an additional 28 antibodies, which included variants predicted to have higher affinity than the wild type antibody in each library (18 total) and negative controls predicted to have similar or lower affinity as the wild type antibody in each library (10 total). Antibodies were produced in Expi293 cells (**Methods**). After confirming expression of each antibody, we next performed surface plasmon resonance to quantify the affinity of each of the 32 antibodies for their target. Consistent with our model predictions, all antibodies predicted by AbMAP to have lower affinity than the wild type antibody did. We furthermore observed that for three of the libraries, antibodies designed using AbMAP to have higher affinity than the wild type antibody did (Figure 4; Supp. Note 1). For the fourth library, we found that antibodies designed by AbMAP showed an improved association constant, a critical feature in therapeutic antibody efficacy (Figure 8; Table 4). Overall, AbMAP had a hit rate of 82% in predicting both strong and weak binders.

**Figure 4:**
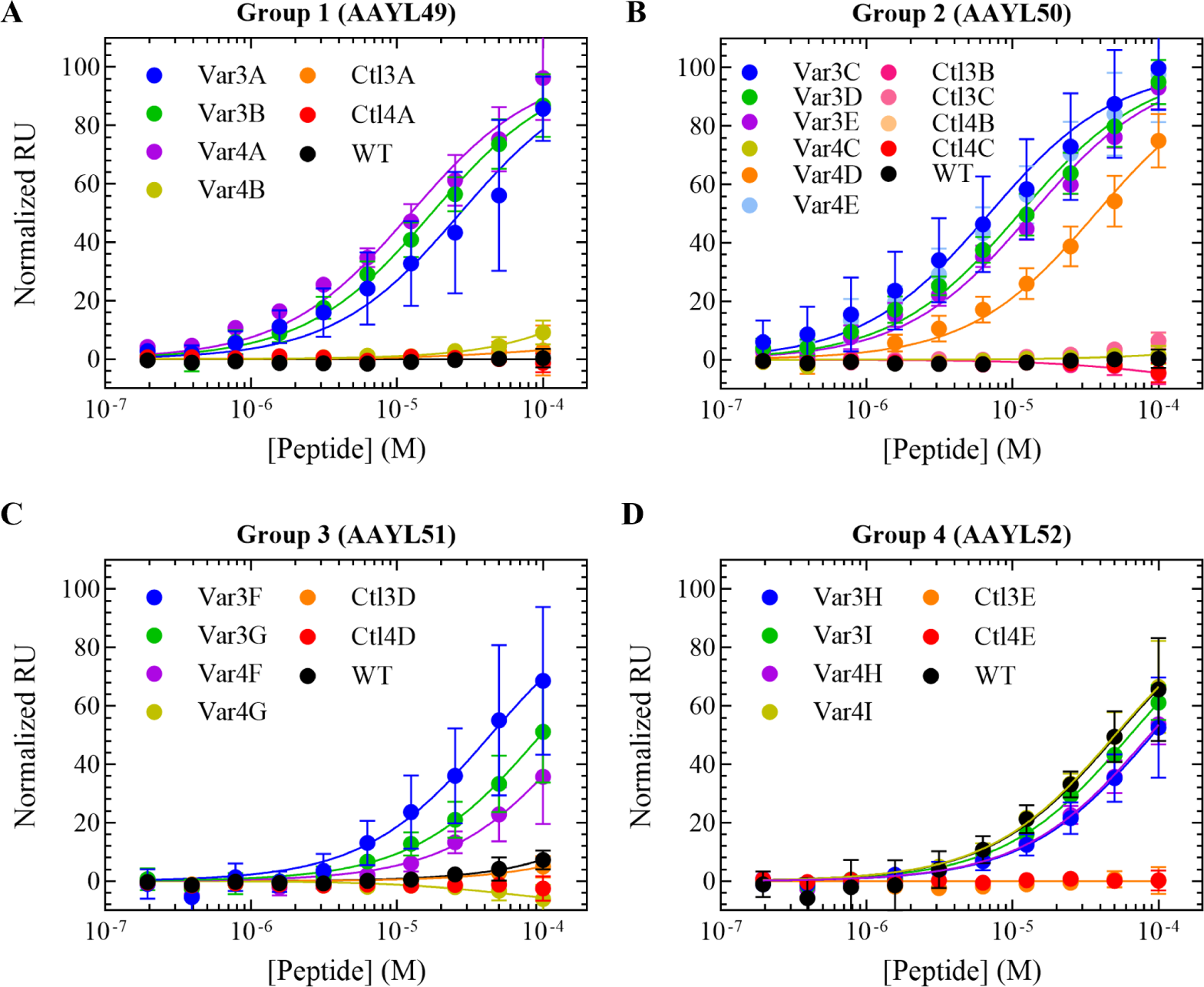
Experimental validation of AbMAP-assessed mutations for antibodies binding to peptide from SARS-CoV-2 spike protein’s HR2 domain. Wild-type sequences from four groups (i.e., libraries) assayed by Engelhart et al. (*19*) were considered. Trained on data from primarily 1- or 2-point mutations in the original study, we predicted and assessed candidate antibodies with 3- or 4-point mutations. Group 2 (AAYL50) was covered more extensively as it contained the best-performing mutants in the original study. Here, normalized binding response (RU) as a function of peptide concentration for the four groups are shown. “Var” and “Ctl” refer to predicted variant and negative controls, respectively, with the number indicating mutations from wild-type (WT). Thus, *Var4A* is a variant with 4 mutations. In all cases, variants were identified with increased or comparable affinity to the WT controls (black). In some instances, the steady state peptide binding affinities approached low single digit nanomolar concentrations. The negative controls showed no binding to the peptide, as expected.

**Table 3:**
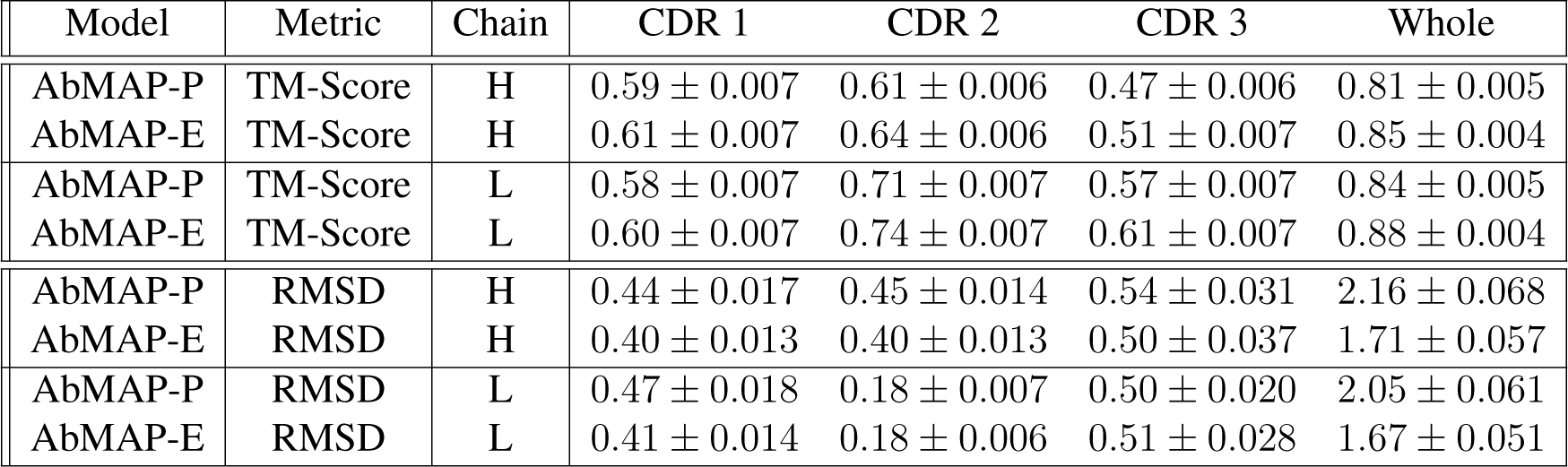
Structure prediction scores for AbMAP-P and AbMAP-E. Similar to AbMAP-B, ProtBert, and ESM-1b in Fig. 1, the predicted template structure was selected from a set of antibodies whose sequence identity is below 0.7. Structure prediction was conducted on both individual CDR fragments and the whole chain.

| Model | Metric | Chain | CDR 1 | CDR 2 | CDR 3 | Whole |
| --- | --- | --- | --- | --- | --- | --- |
| AbMAP-P | TM-Score | H | $0.59 \pm 0.007$ | $0.61 \pm 0.006$ | $0.47 \pm 0.006$ | $0.81 \pm 0.005$ |
| AbMAP-E | TM-Score | H | $0.61 \pm 0.007$ | $0.64 \pm 0.006$ | $0.51 \pm 0.007$ | $0.85 \pm 0.004$ |
| AbMAP-P | TM-Score | L | $0.58 \pm 0.007$ | $0.71 \pm 0.007$ | $0.57 \pm 0.007$ | $0.84 \pm 0.005$ |
| AbMAP-E | TM-Score | L | $0.60 \pm 0.007$ | $0.74 \pm 0.007$ | $0.61 \pm 0.007$ | $0.88 \pm 0.004$ |
| AbMAP-P | RMSD | H | $0.44 \pm 0.017$ | $0.45 \pm 0.014$ | $0.54 \pm 0.031$ | $2.16 \pm 0.068$ |
| AbMAP-E | RMSD | H | $0.40 \pm 0.013$ | $0.40 \pm 0.013$ | $0.50 \pm 0.037$ | $1.71 \pm 0.057$ |
| AbMAP-P | RMSD | L | $0.47 \pm 0.018$ | $0.18 \pm 0.007$ | $0.50 \pm 0.020$ | $2.05 \pm 0.061$ |
| AbMAP-E | RMSD | L | $0.41 \pm 0.014$ | $0.18 \pm 0.006$ | $0.51 \pm 0.028$ | $1.67 \pm 0.051$ |

**Table 4:** Kinetic parameters of the different variants. The affinities (*µ*M) of each variant to the peptide are reported as the average and standard deviation of the steady state model for the indicated number of independent replicates (N). The Rmax indicates the maximum signal of the binding interaction and was used for normalization purposed for direct comparison between the variants. The fold change versus WT is reported for each variant compared to the WT in the appropriate group. If no binding of the appropriate WT was observed (N.B.), the value reported is in comparison to the highest concentration of antigen used in the assay (200 *µ*M) and underestimates the fold difference. N.B. = no binding observed at 200 *µ*M or less of peptide, N/A = not applicable as no binding is observed.

| | | Affinity<br>± SD<br>(μM) | Fold<br>Change<br>vs. WT | Rmax<br>(RU) | N |
| --- | --- | --- | --- | --- | --- |
| Group 1 | Var3A | 18.6 ± 7.8 | >11.8 | 56.6 | 5 |
|  | Var3B | 18.5 ± 5.7 | >10.8 | 39.5 | 6 |
|  | Var4A | 11.6 ± 2.9 | >17.2 | 77.2 | 6 |
|  | Var4B | 74.5 | >2.7 | 18.6 | 1 |
|  | WT | N.B. | -- | N/A | N/A |
| Group 2 | Var3C | 9.5 ± 4.2 | >21 | 50.9 | 6 |
|  | Var3D | 11.9 ± 3.0 | >17 | 63.1 | 6 |
|  | Var3E | 15.1 ± 8.4 | >13 | 43.3 | 6 |
|  | Var4C | 98 ± 33 | >2 | 13.0 | 2 |
|  | Var4D | 39 ± 10 | >5 | 56.7 | 6 |
|  | Var4E | 9.2 ± 4.6 | >22 | 48.8 | 6 |
|  | WT | N.B. | -- | N/A | N/A |
| Group 3 | Var3F | 58 ± 40 | 2.7 | 24.2 | 6 |
|  | Var3G | 113 ± 69 | 1.4 | 51.5 | 6 |
|  | Var4F | 162 ± 12 | 1.0 | 42.9 | 2 |
|  | Var4G | N.B. | -- | N/A | N/A |
|  | WT | 155 | -- | 15.6 | 1 |
| Group 4 | Var3H | 94 ± 36 | 0.7 | 56.6 | 6 |
|  | Var3I | 65 ± 12 | 0.9 | 52.3 | 6 |
|  | Var4H | 91 ± 29 | 0.7 | 66.4 | 6 |
|  | Var4I | 52 ± 20 | 1.2 | 52.2 | 6 |
|  | WT | 61 ± 46 | -- | 39.6 | 6 |

## AbMAP Reveals Shared Landscapes Across Antibody Repertoires

The scalability of our approach, along with its fidelity in capturing structure and function, enables systematic analyses of large antibody repertoires. Existing approaches are ill-suited to such analyses: while structure-prediction methods can not scale to large repertoires, purely sequence similarity-based analyses (*10*) will not be sensitive to structural/functional similarities between antibodies with different sequences. While PLMs directly address this concern, language models that learn the full antibody’s representation (e.g., AntiBERTa (*17*)) may misemphasize the framework diversity at the expense of CDR diversity, with the latter being the key determinants of antibody specificity.

## LIBRA-Seq Cell Line Identification

As a preliminary evaluation, we assessed AbMAP on Setliff et al.’s LIBRA-seq study, in which they profiled B-cell receptor (BCR) binding specificity against a panel of HIV and influenza-related antigens. We wondered whether AbMAP could recover the cell of origin of the BCR. While we had used a subset of this dataset in AbMAP’s multi-task training, we note that no cellof-origin information was provided during training. Furthermore, we additionally evaluated on a subset of BCRs held out during training.

For consistent visualization across analyses, we first standardized a 2-dimensional representation of the overall antibody space. Using the sketching algorithm Hopper (*35*), we selected 1,000 diverse antibodies from the SABDab training set. The farthest-first sampling approach of Hopper is motivated by the vertex k-center problem, and allows us to identify a set of antibodies that cover the entire space. We then computed a 2D reduction by mapping the fixed-length AbMAP-B representations of these antibodies to the top two principal components. In all analyses that follow, AbMAP fixed-length embeddings are reduced to 2D space by this specific mapping, enabling us to visualize different datasets (e.g., in Figures 5a-c) on the same axes. In most plots, we mark the 1,000 “anchor” antibodies as gray dots in the background, to help contextualize the visualization.

**Figure 5:**
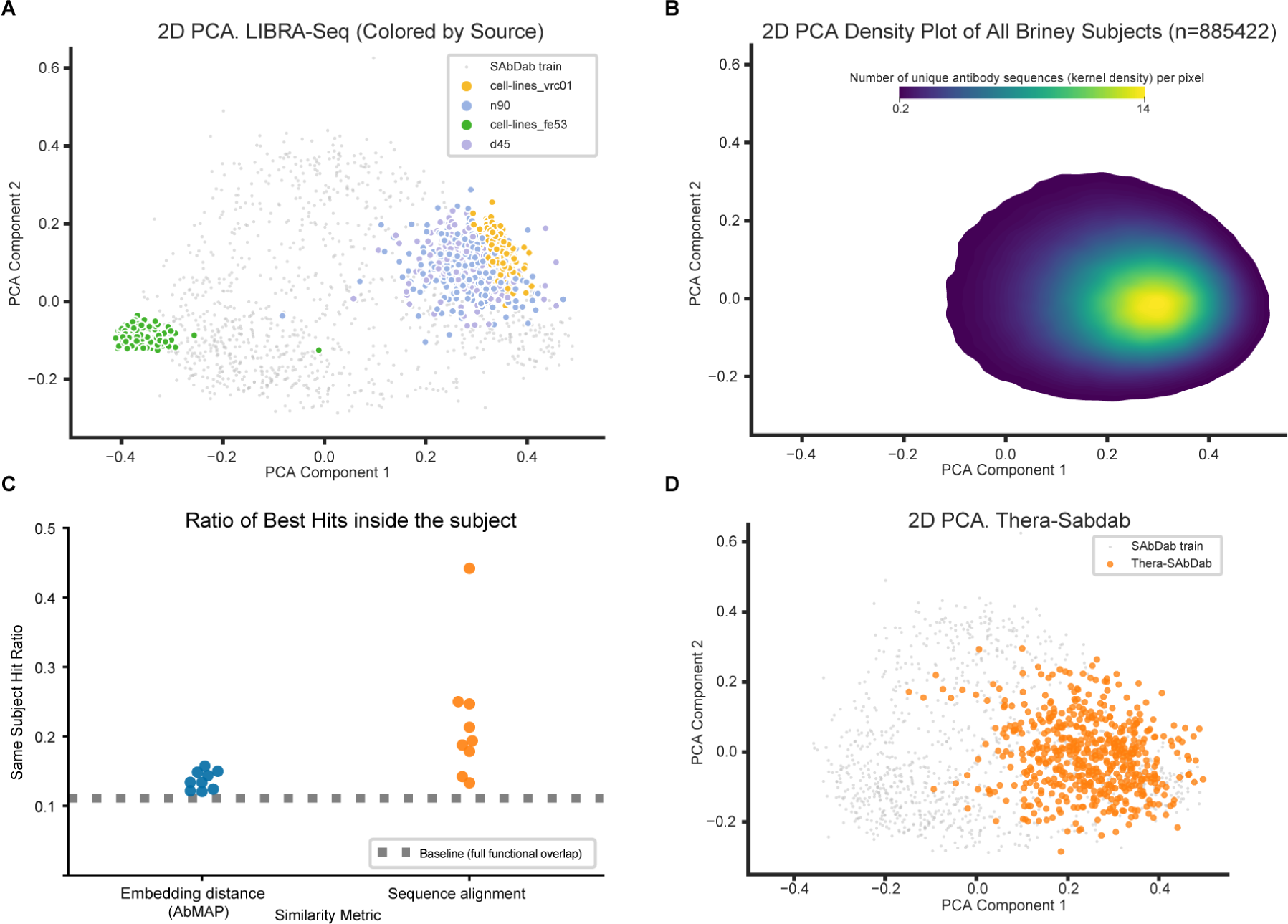
AbMAP embeddings capture structure-function convergence. **A**) PCA of LIBRA-Seq dataset, showing four mini-repertoires. The light gray points represent embeddings of antibodies used during AbMAP training; these were used to compute the PCA. VRC01 is a cell-line derived from a CD4-binding-site-directed HIV-1 broadly-neutralizing antibody (bNAb) while d45 and n90 are from the immune repertoires of HIV-resistant donors NIAID45 and NIAID N90, respectively. Fe53 is a cell-line derived from a group 1 influenza hemagglutinins bNAb. While the HIV-related repertoires are closer to each other than the influenza-related repertoire, the cell line-based populations display less diversity than patient-derived repertoires. **B**) Kernel-density estimate plot of 2D PCA for fixed-length embeddings of antibody repertoire sequences sampled from 9 human subjects sampled by Briney et al. As shown in Fig. 10, there is substantial overlap across per-individual repertoires. **C**) To evaluate if AbMAP embeddings expose structure-function convergence, we measured the overlap across individuals using AbMAP embeddings, comparing it to a sequence-alignment based measure. Each datapoint shows, for one individual, the ratio of sequences where the closest sequence was from the same individual (baseline probability=0.11 when each individual repertoire is an independent sample from an identical distribution). Sequence alignment shows less overlap between subjects than AbMAP embeddings. **D**) 2D PCA plot of 547 antibodies from Thera-SAbDab. We observe that therapeutic antibodies have similar representation as seen in the hotspot of human repertoires (panel B).

We computed AbMAP-B embeddings for 887 LIBRA-seq BCR sequences that were not part of the training set. As shown in Fig. 5a, our model representation was able to differentiate whether BCRs originated from engineered B-cell lines or real human donors. Furthermore, our model was able to discern the two different types of BCRs within the engineered cell lines: VRC01, a CD4 binding-site-directed HIV-1 bNAb (broadly neutralizing antibody), and Fe53, a bNAb recognizing the stem of group-1 influenza hemagglutinins (Fig. 5a). We note that there is some clonal diversity within each cell line, with cells in the line sharing the same lineage but having diversified through antigen exposure and somatic mutation (*36*).

## The Landscape of Human Antibody Repertoires

We analyzed the human antibody datasets from the Briney et al.’s study of BCR repertoires across multiple individuals. As part of this analysis, we ask two key questions:

- Is the set of BCRs uniformly distributed over the embedding space, or does the distribution display “hotspots” of clustering? In case the latter is true, we also wondered if antibody drug candidates that have successfully passed pre-clinical evaluations were more likely to cluster in these hotspots.
- While an extraordinary sequence diversity of BCRs has been reported across individual repertoires (*10*), multiple analyses have suggested that these diverse repertoires converge to similar structure/function (*37, 38, 39*). We wondered if such similarities across individuals would be more evident in our representation space than in the raw sequence space.

Recently, machine learning and experimental approaches have suggested a convergence of structure and function across human BCR repertoires (*37, 38, 39*). Additionally, Friedensohn et al. reported a deep learning approach that shows extensive convergent selection in antibody repertoires of mice for a range of protein antigens and immunization conditions (*40*). AbMAP-based embeddings enable us to go beyond previous approaches in providing a large-scale, principled approach to systematically quantifying the structural/functional convergence across individual repertoires. While previous work has primarily been focused on convergence in sequence fragments, our embedding-based approach encompasses convergence also in “paratope structural signatures” (*39*). We therefore hypothesized that incorporating AbMAP representations in repertoire analysis could more accurately reveal the extent of structural/functional convergence across individuals.

We acquired 100,000 randomly-chosen BCR sequences from 9 individuals each, filtering out identical sequences from an individual. Because of assay limitations in the original study, many of the sequences were truncated on one or both ends. This would be a challenge for language models that need to consider the full antibody; however, our focus on just the CDRs enabled us to recover embeddings for the vast majority (*n* = 885, 422) of these BCRs. We applied AbMAP-B on these sequences and visualized their 2D reduction as described above. We found the distribution to be highly clustered, with a kernel density estimator of the distribution being unimodal (Fig. 5b). Notably, the human cell lines from the LIBRA-seq data overlap very well with the Briney dataset. Interestingly, while the VRC01 cell line falls within the high-occupancy region, the Fe53 cell line is well out of it. While both cell lines are engineered human B cell lines (Ramos) (*41*), they express distinct BCRs that neutralize different antigens (VRC01 antibody neutralizes HIV-1 while Fe53 antibody neutralizes influenza).

We next assessed if the antibody repertoires across individuals were similar. In the original study, Briney et al. had observed substantial cross-individual diversity. While this makes sense given the vast space of possible antibody sequences, it is also somewhat puzzling— ultimately, most individuals need antibody-based protection from similar antigens (e.g., the flu, environmental stressors etc.). Indeed, when we visualized each individual’s repertoire in our embedding space, we found that the distributions looked remarkably similar (Fig. 10), suggesting that each repertoire had similar structural/functional coverage. For a more systematic assessment of cross-individual overlaps, we sampled 5,000 sequences from each subject (i.e., 45,000 sequences in total) and performed all-vs.-all pairwise comparisons, using either the raw sequence^1^ or the AbMAP-B representation (**Methods**). From these pairwise comparisons, we obtained the nearest neighbor of each antibody across all individuals and computed the frequency with which this neighbor hails from the same individual. If the per-individual repertoires are independent and identically distributed (i.i.d.) samples from the same underlying distribution, the fraction of cases where the nearest neighbor is from the same individual should be 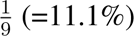. When using AbMAP-B embeddings to compute similarities, we found that this to nearly be the case (Fig. 5c): per-individual fractions averaged 0.14 *±* 0.013, with very little variation across individuals. In contrast, when using the sequence similarity metric, the average per-individual fraction was substantially higher (0.22 *±* 0.087), with much greater variability across individuals. Thus, while per-individual repertoires seem to differ substantially when only sequence similarity is concerned, these repertoires are revealed to be much more similar in their structure/function coverage when represented by AbMAP.

We wondered if the distribution pattern of human antibody representations is therapeutically useful. Towards that, we mapped 547 antibodies from Thera-SABDab, a dataset of immunotherapeutic antibodies that have entered clinical trials (*42*). Our hypothesis was that even if an antibody drug candidate is effective in vitro, it may not have drug-like properties (e.g., low toxicity) unless it falls solidly within the realm of native-like antibodies. As Fig. 5d indicates, this does seem to be the case – the set of antibodies in Thera-SABDab are located in the high-occupancy region of the embedding space. Thus, a general drug-discovery recommendation we make is to sample candidate antibody drugs primarily from this region of the embedding space.

## Predicting SARS-CoV-2 Variant Neutralization Ability

We also examined AbMAP’s capability on the prediction of antibody efficacy on neutralizing SARS-CoV-2 variants. This analysis also demonstrates how AbMAP could be applied in a low-N setting to optimize an existing panel of antibodies to address incremental mutations in the target. From CoV-AbDab, a dataset of coronavirus-binding antibodies, we obtained the set of 2,077 antibodies that were reported to neutralize the wild-type strain of SARS-CoV-2; we then computed fixed-length AbMAP-B embeddings for these. Setting aside 522 (approx 25%) randomly chosen antibodies, we plotted the rest (=1,555) on the 2D visualization described above (Fig. 6a). The set of wild-type neutralizing antibodies span a fairly large part of the overall space, suggesting that a substantial diversity of antibody structures is capable of neutralizing SARS-CoV-2 (we note that these antibodies vary in their reported viral target protein).

**Figure 6:**
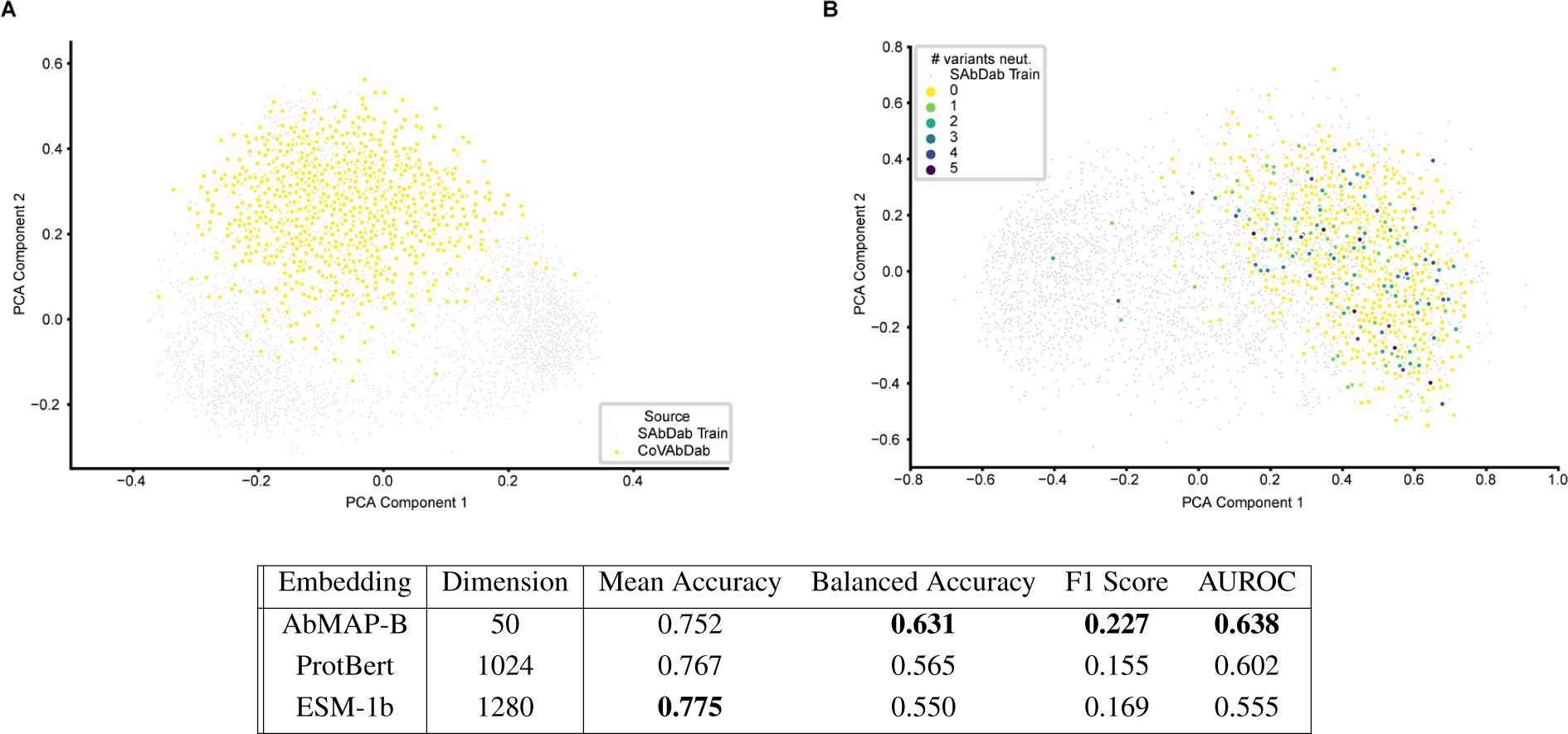
Applying AbMAP to predict SARS-CoV-2 variant neutralization. On the 2-D PCA computed in Fig. 5 (SAbDab training data in gray), (A) we mapped SARS-CoV-2 neutralizing antibodies (sourced from CoVAbDab). (B) We fitted a linear projection of AbMAP embedding that distinguishes CoVAbDab antibodies from other antibodies. No variant-neutralization information was used to compute this projection. We then assessed if the learned AbMAP-informed CoVAbDab representation could predict variant neutralization efficacy. We measured efficacy as the ability to neutralize at least one variant, and used 5-fold cross-validation to benchmark AbMAP against baseline PLM models.

Seeking to further hone in on the variant neutralization capabilities of these antibodies, we applied Schema (*43*) to extract a linear projection of AbMAP-B embeddings that accentuates the difference between CoV-AbDab antibodies and a baseline set of 3,000 antibodies (the SAbDab subset used to train AbMAP). Schema computes a low-distortion linear projection of an embedding such that distances in the projected space better capture a user-specified co-variate; here, membership in CoV-AbDab (Fig. 6b). By applying Schema, we are able to enrich the SARS-CoV-2 relevant signal in the embedding so that a simple model, working with relatively limited training data, can capture the desired biological intuition. On the held-out set of 522 CoV-AbDab antibodies, we annotated each antibody with a binary label indicating if it neutralizes at least one SARS-CoV-2 variant (Alpha, Beta, Delta, Gamma, Omicron). We then applied logistic regression to predict this label from the Schema-projected representation. We evaluated the results as per 5-fold cross validation, also comparing fixed-length embeddings derived from ProtBert and ESM-1b (Fig. 6). AbMAP-based representation, even though it is of much lower dimensionality (50 compared to ESM-1b’s 1,280) has substantially higher balanced-accuracy and F1 score, indicating that our transfer learning approach is more effectively able to hone in on the subspace relevant to SARS-CoV-2 neutralization.

## Discussion

We presented AbMAP, a transfer learning framework to adapt any foundational PLM (i.e., one trained on the broad corpus of protein sequences) to antibodies, whose hypervariable regions lack the evolutionary conservation that PLMs typically rely on. Consider a foundational PLM’s representation of a residue in a complementarity-determining region (CDR): it captures information about the residue in the context of the overall protein. However, this context was learned from the distributional properties of general proteins while the hypervariability in a CDR implies a distinct distributional context. We therefore refine the raw PLM embedding by *in silico* mutation-based contrastive augmentation of the CDR residues. To maximally leverage the limited structural data on antibodies, AbMAP also focuses its capacity on the CDRs, these being the key drivers of antibody specificity. As CDR signatures in antibody sequences have been well characterized, a hidden markov model like ANARCI can reliably identify such regions in an antibody sequence (*22, 23*). In comparison, a model seeking to learn the full antibody sequence (e.g., AntiBERTa (*17*)) may expend significant portion of its learning capacity on re-discovering such signatures during training. To mitigate the trade-off in focusing on hypervariable regions rather than the full antibody, we include two flanking residues on each side of a CDR to partially capture the role of framework residues.

To further shape the augmented embedding, we supervise its non-linear projection to a lower dimensional space such that Euclidean distances in the projected space better capture antibody structural and functional similarity. In an ablation study, we found that training on just structural data recapitulated function but not vice-versa (Table 5). This may be due to single-cell data sparsity and noise in the LIBRA-seq dataset (*26*) we used for the latter. Correspondingly, our multi-task formulation also learned a higher weight for the structural task. We present AbMAP-E, AbMAP-P, and AbMAP-B, adaptations of the **E**SM-1b, **P**rotBert, and the **B**epler & Berger PLMs, respectively. The choice of foundational PLM may depend on the type of downstream prediction task using the resulting AbMAP embedding (e.g., we found AbMAP-B to be better for structure prediction, and AbMAP-E/P for function/property prediction). Our transfer learning framework can be adapted to any new PLM, and we make both the training data and code available for doing so.

**Table 5:** Test set Spearman’s rank scores computed for AbMAP-B,E when using different datasets for train and test (i.e. we evaluated the prediction of functional similarity using embeddings trained on structure data, and vice versa). While embeddings trained solely on structure data perform quite well when predicting function (underlined), embeddings trained only on function data can infer far less about antibody structure.

| Model | Train Data | Test Data | Spearman's $\rho$ |
| --- | --- | --- | --- |
| AbMAP-B | SAbDab | LIBRA-Seq | <u>0.57</u> |
| AbMAP-B | LIBRA-Seq | SAbDab | 0.028 |
| AbMAP-E | SAbDab | LIBRA-Seq | <u>0.60</u> |
| AbMAP-E | LIBRA-Seq | SAbDab | 0.049 |

AbMAP represents a conceptual advance to language model design for antibodies. Currently, two broad approaches have been espoused. One, embodied in techniques like OmegaFold (*6*), is to essentially focus on improving the general-purpose PLM, with the expectation that gains will accrue to antibodies as well. The other, represented by methods such as AntiBERTa (*17*), AbLang (*44*), and IgLM (*18*), is to essentially treat antibodies as an entirely new language-modeling task and train solely on corpora of antibody sequences. AbMAP represents a new middle path. We describe a transfer learning approach that can adapt to any foundational PLM and hence benefit from innovations in the underlying PLMs. Since antibody hypervariable regions are not evolutionarily conserved, foundational PLMs will likely remain weaker at modeling antibodies than regular proteins. Our approach starts from informative foundational PLMs and then adapts them to be more accurate on antibody structure and function. We additionally make the choice to focus AbMAP’s explanatory capacity on the CDRs and their flanking residues. This is an inductive bias, allowing us to limit the model complexity of the transfer learning layer (one layer of transformer) and enabling robustness as well as accuracy. Lastly, an intriguing direction of future work would be to combine ideas from AbMAP’s transfer learning approach with repertoire-based PLMs (like AntiBERTa) to infuse structural, functional as well as repertoire information into antibody representations.

We demonstrated AbMAP to be applicable in a variety of cases, e.g., property prediction and antigen binding. We briefly elaborate on AbMAP’s strengths in the context of structure prediction, as this is an area of active research and there exist methods focused solely on antibody structure prediction. Our implementation of structure prediction with AbMAP is as a template-finding task, and we leave the task of template refinement to future work. Even unrefined, however, AbMAP performs remarkably well, outperforming AlphaFold 2 and being competitive with OmegaFold in most cases. It outperforms the latter on the functionally-crucial CDR-H3 region. We wondered if our template-finding approach was biasing the results in AbMAP’s favor, and re-evaluated it at progressively lower levels of sequence homology (Fig. 7); we also compared AbMAP-*{*B,E,P*}* with their respective foundational PLM baselines. AbMAP’s performance did not decline meaningfully at lower sequence homologies and it substantially outperformed its foundational PLM baselines. Notably, the latter did not compare as favorably with OmegaFold or AlphaFold2, suggesting that it is our transfer learning innovations, rather than the template-based approach, that offers the gains.

**Figure 7:**
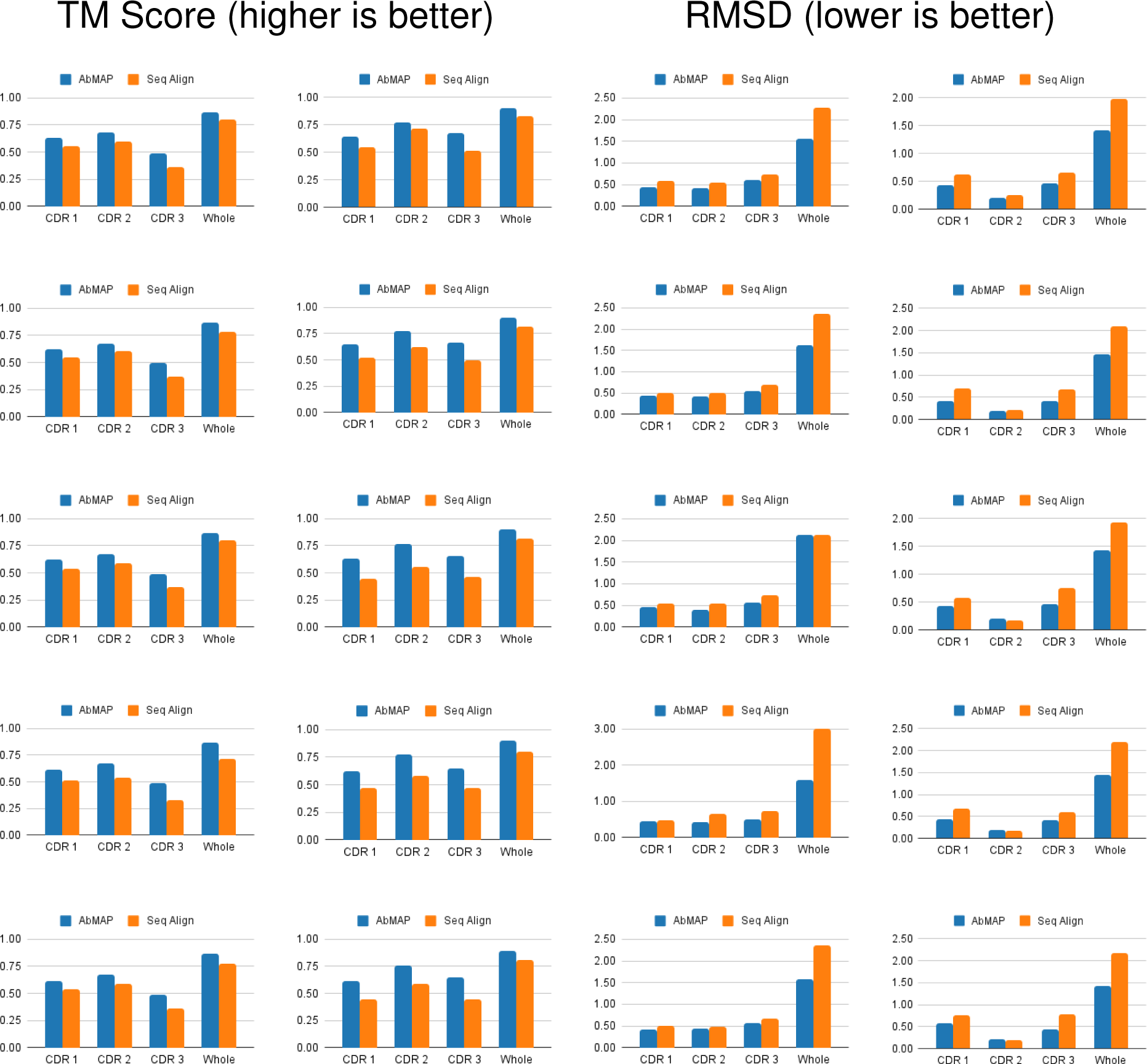
Comparison of AbMAP-B and sequence alignment at different sequence identity thresholds for template search in antibody structure prediction. Columns from left to right (TM-Score Chain H, TM-Score Chain L, RMSD Chain H, RMSD Chain L). Rows from top to bottom (Sequence identity: 0.9, 0.8, 0.7, 0.6, 0.5). For a pair of antibodies, AbMAP-B uses the cosine distance of fixed length sequence embeddings and sequence alignment uses the pairwise global sequence alignment score as a similarity metric.

**Figure 8:**
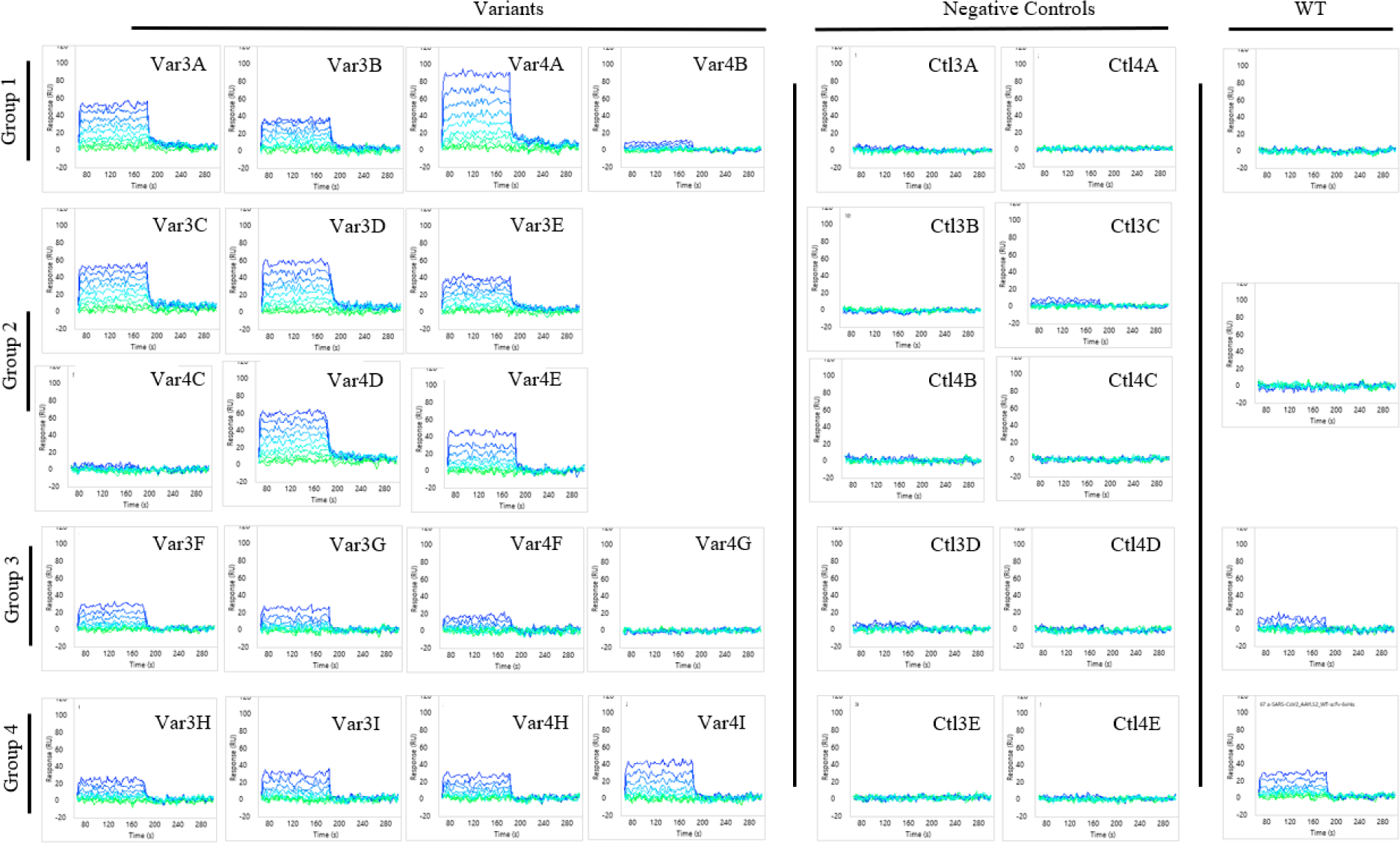
Concentration dependent binding of the SARS-CoV-2 peptide to each scFv antibody. Sensorgrams are organized by group with the appropriate negative control and WT sequence.

**Figure 9:**
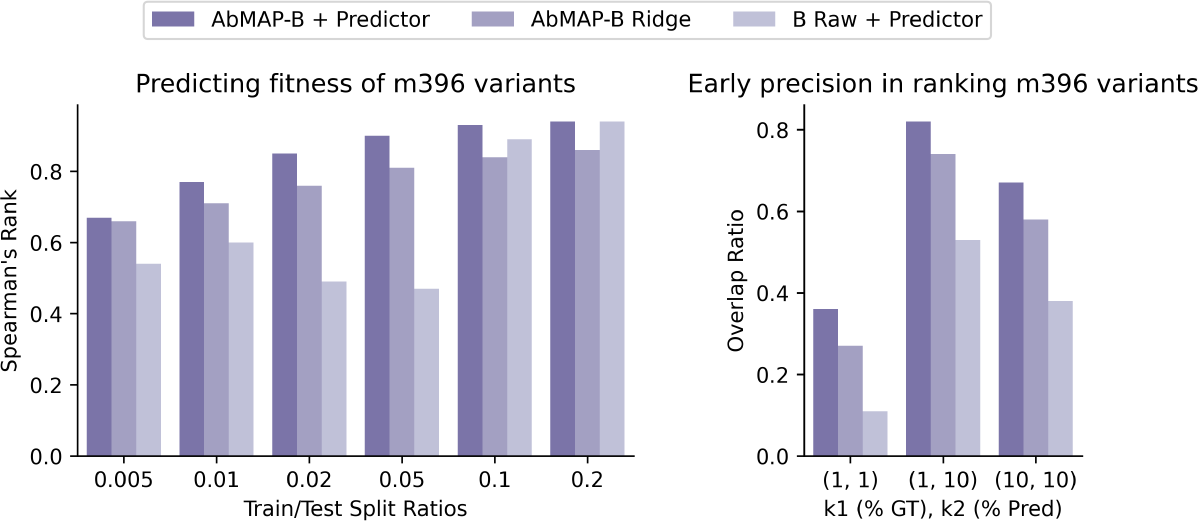
(Left) Chart of average Spearman’s rank scores for ddG scores prediction of Bepler & Berger based models (AbMAP and raw) with various train/test splits. (Right) Chart of average overlap of top-*k*_1_, *k*_2_ ddG scores for different *k*_1_, *k*_2_ values. This experiment was conducted for different train/test splits (train: 0.02, test: 0.98 for this chart). While the raw embeddings are comparable to augmented embeddings at higher splits, their performance notably drops at lower training set sizes.

**Figure 10:**
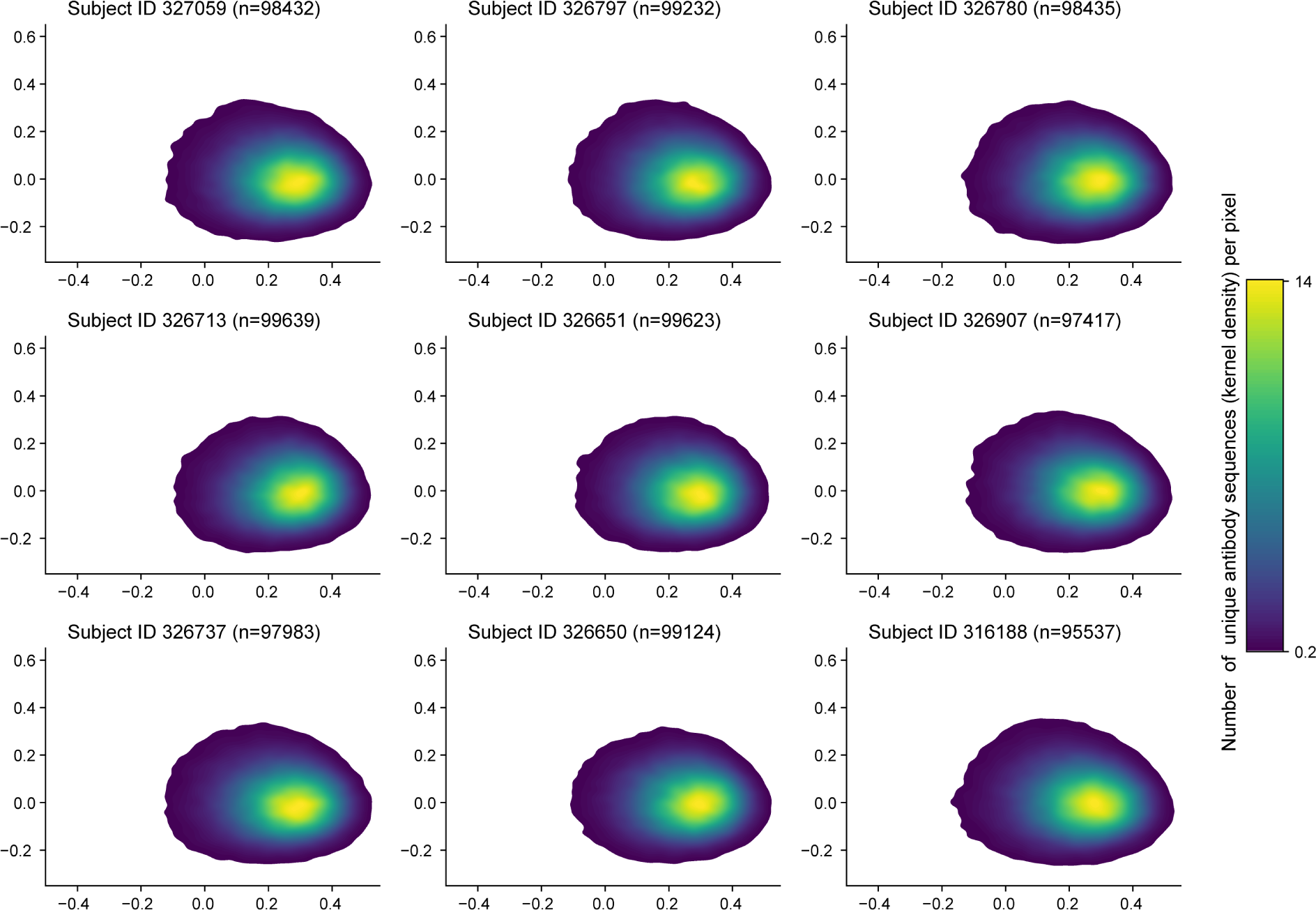
KDE plot: 2D PCA of AbMAP embeddings for antibody repertoire from each human subject.

AbMAP’s design is compatible with a variety of foundational PLM architectures. Here the three foundational models we use differ substantially in their approach: the Bepler & Berger model was trained in a multi-task setting and also uses protein structure information while the ProtBert model adapts the existing BERT architecture, and the ESM-1b model uses a novel transformer-based architecture. Across a variety of applications, the corresponding AbMAP variants outperform their foundational baselines. We believe this is due to our focus on a) CDRs, b) contrastive augmentation that accentuates the residues in the CDRs, and c) a modular architecture with the fine-tuning layer cleanly separated from the foundational PLM. Of these, we believe the last to play the most important role (Fig. 2). In our fine-tuning layer, we apply a single transformer layer to adapt contrastively-augmented representations into final AbMAP embeddings; we found that additional layers did not meaningfully increase performance in terms of test set loss and Spearman’s rank (Table 6).

**Table 6:** Test set mean loss and Spearman’s rank scores computed for AbMAP-B when using different number of Transformer Encoder layers in its architecture. Increase in number of layers did not necessarily show improvement in model performance.

| num. transformer layers | 1 | 2 | 3 |
| --- | --- | --- | --- |
| Mean Loss | 0.272 | 0.281 | 0.279 |
| Spearman's $\rho$ | 0.826 | 0.805 | 0.817 |

We demonstrated AbMAP’s applicability to antibody design, optimizing a panel of SARS-CoV-2 antibody candidates. Our predictive framework offers numerous advantages. It enables the generation and evaluation of sequences of arbitrary lengths. Unlike diffusion-based generative approaches optimized solely for finding strong binders, our framework can also efficiently generate negative controls. The clustering-based candidate selection offers a tunable balance between exploration and exploitation: in early rounds, a wider range of clusters can be explored, while in later stages, specific clusters may be focused on. This approach scales to consider millions of candidates, a volume unmanageable for Bayesian optimization using Gaussian processes.

In contrast, existing in silico approaches have generally reported substantially lower hit rates. The recent version of RFdiffusion, a generative binder-generation approach for antibody, has reported hit rates of under 5% (*45, 46*). A language model-based approach to antibody optimization by Bachas et al. reported a 74% hit-rate for binders that achieved the desired objective (*47*). Notably, Bachas et al. did not use a foundation model and sought to learn representations solely from antibody data; this suggests that our transfer learning approach can offer advantage especially in situations with limited data. Taken together, the results of this validation study suggest that AbMAP ’s efficiency in identifying strong binders expedites the discovery and optimization of antibody therapeutics, especially against rapidly evolving pathogens such as SARS-CoV-2.

The design of AbMAP implies some trade-offs. Our explicit focus on the hypervariable region of the antibody is data efficient and offers us robustness to immunoglobulin isotype switching during repertoire analyses. However, it may come at the expense of capturing the role that the framework region may play in an antibody’s stability and specificity. To mitigate this, we followed Liberis et al. (*24*) and expanded the standard Chothia delineation of a CDR to also include two flanking residues on each end. In future, as more antibody datasets become available (e.g., those measuring binding or functional specificity in diverse contexts), the additional training data may enable us to leverage more complex models that effectively capture both the framework and hypervariable regions. Also, while our contrastive augmentation step improves performance and leads to more stable results, it also requires multiple invocations of the baseline PLM (one for each mutant). In situations where speed is crucial, this step can be removed for greater efficiency. The model without contrastive augmentation is faster during training and inference, though at the expense of lower consistency with ground truth structural properties (Fig. 11).

**Figure 11:**
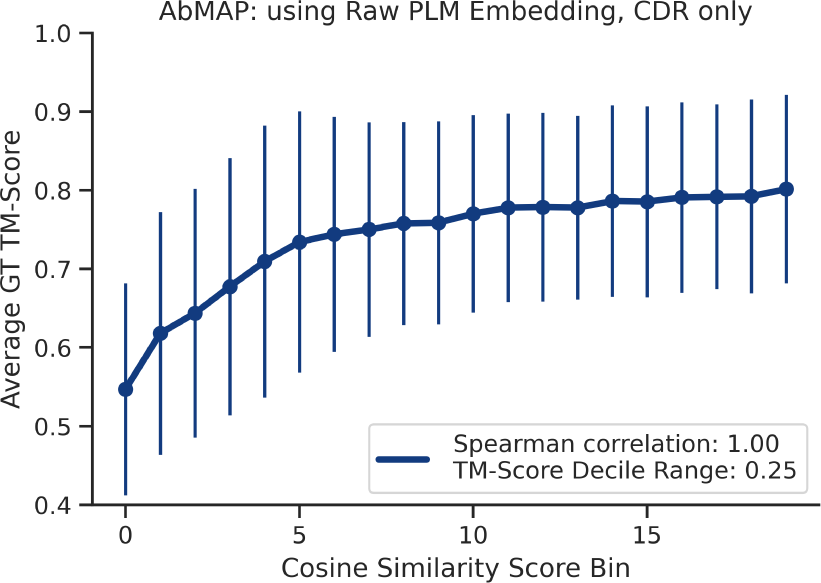
Average ground truth TM-scores antibody pairs in each of the 20 cosine similarity score bins using AbMAP representations with raw PLM embeddings (CDRs only) as input; i.e. no contrastive augmentation. Monotonicity is noticeably lower when using raw PLM embeddings as input to AbMAP compared to using contrastively augmented PLM embeddings.

The exploration of human immune repertoires, as well as the design and development of large-molecule therapeutics, require a deep understanding of antibody structure and function, and an ability to efficiently manipulate it in silico. However, the hypervariability of antibodies poses a challenge for general-purpose protein modeling techniques, hindering their translation to antibodies. The transfer learning approach of AbMAP represents a broadly-applicable solution to such conundrums. Rather than training a dedicated language model from scratch for a sub-category of proteins, we argue it is more effective to leverage the fast-moving advances in foundational PLMs and fine-tune a general PLM towards the subset. AbMAP’s transfer learning framework enables us to effectively adapt foundational PLMs. We believe the scalable and accurate modeling of antibodies will unlock a better understanding of the behavior of antibodies, empowering the discovery of novel therapeutic biologics.

## Acknowledgments

This work was supported by Sanofi, including a research grant from Large Molecule Research and a 2022 iDEA award, as well as the Abdul Latif Jameel Clinic for Machine Learning in Health.

Conflicts of Interest: T.S., Y.Q., B.M., A.G., M.W. and Y.F.N. are/were employees of Sanofi Inc. The other authors have no competing interests.

## Supplementary materials

### Methods

#### Datasets

##### SAbDab

SAbDab (Structural Antibody Database) (*25*) is a database of antibody structures where each structure is paired with various metadata including the heavy and light chains of antibodies in the PDB complex. Limiting ourselves to PDB entries where both the heavy and light chains were available, we obtained 3,785 pairs of heavy and light chain antibody sequences and PDB structures from SAbDab. We divided them into 3,000 pairs for train and 785 pairs for evaluation. From the 3,000 antibodies selected for training, we randomly sampled 100,000 pairs of antibodies and computed the pairwise similarity scores using their PDB structural data. For each of the sampled 100,000 SAbDab antibody pairs, we computed the TM-Scores of the pairs of antibodies (separately for heavy and light chains), and used them as the ground-truth label when supervised against the predicted similarity score from our model. We used TM-Score as the similarity metric as it is length-independent and is a standard metric used in well-known protein structure prediction tasks such as CASP (*48*).

##### LIBRA-Seq

To train the model concurrently on the functionality of antibodies in addition to their structural properties, we used LIBRA-seq, which maps 4,644 B-cell receptor sequences to their antigen specificity (*26*). The antibody sequences were divided into 3,715 and 929 antibodies for train and evaluation, respectively. The 4,644 antibody sequences, each with a heavy and a light chain, were paired with three scores that indicate the binding specificity to the surface proteins from three different antigens: BG505, CZA97, and H1-A/New Caledonia/20/1999 (*49, 50, 51, 52*).

For each antibody, this set of three scores (each standardized to *µ* = 0*, σ* = 1) serves as its binding specificity vector. For a pair of antibodies in the LIBRA-seq dataset, we scored their functional similarity as the dot-product of their vector representations. From the 3,715 antibodies for training, we randomly sampled 100,000 pairs of antibodies and computed these pairwise similarity scores. Similar to the structural similarity prediction mentioned above, these scores were then used as ground-truth labels for the supervised training of our model’s functional similarity prediction task.

##### CoV-AbDab

CoV-AbDab is a database of all published/patented antibody sequences (7,964 sequences in total as of July 19, 2022) that are capable of binding to coronaviruses including SARS-CoV-2, SARS-CoV-1, and MERS-CoV (*53*). To evaluate our trained model, we selected from these the 2,077 sequences that bind to wild-type SARS-CoV-2. On these, we assessed the model’s ability to predict if the antibody could also neutralize at least one SARS-CoV-2 variant (alpha, beta, omicron, etc.), using only the antibody sequence information.

##### Thera-SAbDab

Thera-SAbDab is a set of antibodies (547 in total) collected from the PDB that contain immunotherapeutic variable domain sequences (*42*). Each of these sequences were labeled with data such as the highest clinical trial passed (Phase-I, Phase-II, etc.). We used our language model to compare the distribution of therapeutic antibodies against the human antibody repertoire.

##### Human Antibody Repertoire

The Briney et al. (2019) dataset contains over 3 billion B-Cell antibody sequence reads across 9 human subjects (*10*). For our experiments, we sampled non-unique 100,000 antibody sequence reads from each subject, where we labelled each unique antibody with the read count within each sample population.

##### Datasets for Labeled Antibody Mutants

The dataset provided in Desautels et al. (2020) was used to validate our antibody language model through ddG score predictions (*33*). They used a machine learning model to search the mutational combinatorial space of the m396 antibody, and calculated the binding propensity of these mutants against the SARS-CoV-2 spike protein’s receptor binding domain. They applied five standard software packages to score the physicochemical characteristics of the mutant’s binding, including STATIUM, FoldX and Rosetta (*32, 30, 31*). To validate our model, we used these pre-computed scores to formulate regression-based property prediction tasks, with only the mutant sequence as input.

#### Embedding Generation via mutational augmentation

Given a heavy or a light chain sequence of length *n* of an antibody, we created its *CDR-specific* embedding by performing the following set of operations. We first create an R*^n×d^* embedding using an foundational protein language model such as the Bepler & Berger (*12*), ESM-1b (*11*), or ProtBert (*14*), where *d* = 2200 (Bepler & Berger), 1280 (ESM-1b), or 1024 (ProtBert). We note that Bepler & Berger has 6165 dimensions across three layers but we used only the last layer’s weights. Then, using ANARCI (*20*), we number the amino acid residues in the input sequence using the Chothia scheme (*21*). Then, we identify parts of the embedding that correspond to the CDRs based on the numbering scheme. Next, we generate *k* (here, *k* = 100) new antibody sequences through *in silico* mutagenesis of the original sequence, where the mutation is performed (sampling from a uniform distribution over all amino acids) with certain probability (here, 0.5) for each residue in the identified CDRs. This procedure is repeated *k* times for the original sequence, generating *k* new sequences which are then embedded using a foundational PLM as in step 1. Then, each of *k* new embeddings is subtracted from the embedding of the original sequence, and these adjusted embeddings are averaged. Finally, we extract and concatenate parts of the averaged difference embedding that belong to the CDRs as determined previously by ANARCI, creating an R*^nt×d^ CDR-specific* embedding. The length *n^t^* usually ranges between 20 to 30.

#### Language Model Refinement with a Multi-task Architecture

Once we generate a CDR-specific embedding for a given antibody sequence, we use it as input to our model which outputs a fixed-length feature that can be used for further downstream task-specific similarity prediction. We curated two datasets for pairs of antibodies, one labeled with their 3D structure similarity, and the other with functional similarity (with regards to antigen binding profiles) to train the model. The overall pipeline as well as the diagram of our antibody language model is shown in Fig. 1. Our model consists of an MLP projection module whose parameters are shared across the training of each task (structure and function). This projection module reduces the dimension of the input embedding (R*^nt×d^ →* R*^nt×dt^, d^t^ < d*) so that the language model is forced to retain as much information about the antibody sequence as possible while trying to execute downstream prediction tasks. For example, when using the Bepler & Berger language model as the foundational PLM, the input CDR-specific embedding is R*^nt×^*^2200^ and our model’s projection module outputs a R*^nt×^*^256^ embedding.

Then, the embedding with reduced dimensions, outputted by the projection module is fed into two separate PyTorch (v1.11.0) Transformer Encoder modules, one each for a downstream similarity score prediction task (structure and function) (*54*). We add sinusoidal positional encodings to the input embeddings in order to inject information about the relative positions of amino acid residues in each sequence. The representations for each residue in the embedding outputted by this Transformer Encoder module now incorporate antibody-related structural and functional information by leveraging attention from other residues in the CDR.

In a multi-task training framework such as our setting, it is important that the calculated losses from each task is combined carefully so that the shared parameters in our model is correctly optimized. Rather than using a set ratio to weigh the two MSE losses, we assigned a new learnable parameter *α* for weighing the losses. The overall MSE loss *L_MSE_*is calculated as a weighted sum of the losses calculated for structural similarity prediction (*L_structure_*) and functional similarity prediction (*L_function_*):

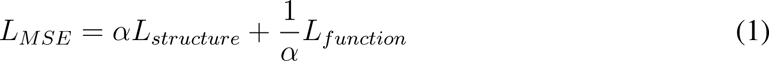

where *α* is updated iteratively through gradient calculations.

#### Fixed-length Embeddings

In addition to the per-residue embeddings (variable length) focusing on the CDRs of an antibody’s sequence, our model can also generate fixed-length embeddings by performing pooling operations on the variable length embedding outputted by the Transformer Encoder along the sequence dimension. Specifically, we use the LogSumExp (LSE) operation, defined as:

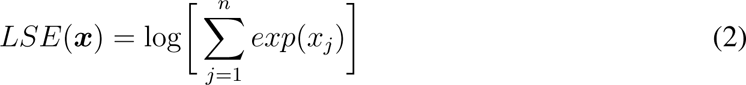

where ***x*** = (*x*_1_*, …, x_n_*), to compute a smooth maximum over the sequence embedding. We also use mean pooling to generate another fixed length embedding of same length. The two are then concatenated into a single representation. Overall, the variable length feature in space R*^nt×dt^* is transformed into a fixed length feature in space R*^q^*, where *q* = 2*d^t^* (here, *q* = 512). This fixed-length vector for each input antibody sequence is then used for similarity score prediction using the cosine distance metric.

#### Max-Entropy regularization

In order to prevent the learned representations being overfit to the datasets used for supervision, we include a regularization loss adapted from Shannon entropy (*55*):

[math3

The regularization is applied to the antibody’s *L*^2^-normalized fixed-length feature of length *q*, which is outputted by the task-specific Transformer Encoder module following the nonlinear projection module. The square of each entry in the fixed-length feature is treated as the probability of an arbitrary discrete random variable *X*, corresponding to *P* (*x_i_*) in the above equation. To induce a regularizing effect, we want the squared entries in the feature to form a uniform distribution. For each *L*^2^-normalized feature, the squared entries are non-negative and sum to 1, like a probability distribution. We therefore set the regularization loss in a max-entropy formulation as:

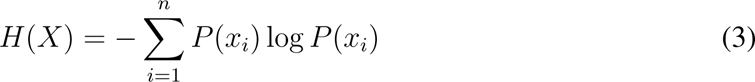

where *u* = [*u*_1_*, u*_2_*, …, u_q_*] is a task specific feature of length *q*.

Therefore, with a regularization parameter *λ*, the total loss computed during a single feedforward step is:

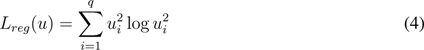

where *λ* was empirically determined as 0.0005, and was used throughout our training.

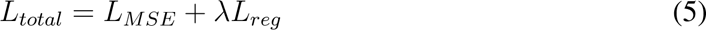

#### Alignment of Raw Antibody Sequences

We used the Needleman-Wunsch algorithm (*56*) to compute the per-residue similarity of two protein (antibody) sequences using the implementation of the algorithm in BioPython (v1.69). We set the match (identical character) and mismatch scores to 1 and 0, respectively, and did not assign gap penalties.

#### Experiment Setup

We trained our model for 50 epochs with a batch size of 32 for each task. We used stochastic gradient descent (SGD) optimizer to update the model parameters. Our multi-task learning model contains approximately 4.5 million parameters. In addition to the regularizations described above, we also included a dropout layer after the projection step. The model was loaded and trained on a single NVIDIA Tesla V100 GPU, and it took around 3.5 hours to train the model.

#### Iterative Antibody Optimization: Experimental Validation

##### Surface plasmon resonance assay for SARS-CoV-2 peptide binding

For the biological validation phase, the variable fragment (Fv) regions of each antibody were reformatted into a scFv containing a C-terminal hexahistidine tag and inserted into a pTT5 mammalian expression vector for production in Expi293 cells. In total, 32 candidate scFvs across the four libraries were proposed, consisting of the 3 original wild-types, 18 positive predictions and 10 control predictions, with 29 (91%) having measurable quantities post-purification (*>* 10*µ*g). Each scFv was assessed for purity and integrity by LabChip prior to subsequent use. To assess the binding affinities of the scFv, we employed Surface Plasmon Resonance imaging (SPRi, Carterra), a method offering high throughput and sensitivity for molecular interaction measurements.

##### Antibody synthesis

Each antibody Fv sequence was reformatted into a scFv and cloned into a pTT5 mammalian expression vector. A 15 amino acid glycine4-serine repeat linker (G4S)3 was inserted between the VH and VL domains to ensure flexibility and a hexahistidine-tag was added at the scFv C-terminal to enable purification. Each scFv was transfected into Expi293 cells according to the manufacturer’s specifications at a DNA ratio of 1 *µ*g DNA per mL of culture. Five days post-feed, the supernatant was isolated and filtered through 0.22 *µ*m filters to remove any aggregates and remaining cells and cell debris. The supernatants were diluted 1:1 with binding buffer (20 mM sodium phosphate, 300 mM sodium chloride, pH 7.4) and flowed at 1 mL/min over 1 mL hisTrap HP columns (Cytiva). The columns were washed with wash buffer (20 mM Sodium Phosphate, 300 mM sodium chloride, 10 mM imidazole, pH 7.4) for a total of 10 column volumes, prior to elution with 5 column volumes of elution buffer (20 mM sodium phosphate, 300 mM sodium chloride, 500 mM imidazole). The proteins were buffer exchanged into PBS, pH 7.2 for subsequent experiments. The molar extinction coefficient for each scFv was used to determine the concentration by UV absorbance at 280 nanometers. Purity was determined by LabChip according to the manufacturer’s instructions and loading 2.5 *µ*g per sample.

### Supplementary Notes, Figures, and Tables

#### Supplementary Note 1: Discussion of iterative antibody optimization

##### Assay differences between our study and Engelhard et al

Our surface plasmon resonance (SPRi) assay differs from the phage display methodology from the original study. Phage display technologies display a valency effect which may confound the apparent affinity measurements (e.g., kD). In contrast, the SPRi assay offers two key advantages: a) it eliminates the need to scout for capture conditions to obtain significant immobilization levels for the binding study; and b) steady-state measurements can be obtained even for weak interactions. Here we assayed the normalized binding response (RU) as a function of peptide concentration for groups. Coupling of the scFv to the chip surface in our assay was also valuable as the relatively small small size of the antigen peptide (¡3 kDa) compared to intact protein domains further necessitated a direct immobilization scheme to ensure appropriate signal to noise (¿10 RU).

##### Analysis of results

The series of concentrations of the antigen was injected over all the immobilized antibodies simultaneously and a majority (¿80%) displayed some level of binding signal (5 RU) at the highest antigen concentration tested in the assay (100 *µ*M). Figure 8 highlights the sensorgrams for all molecules based on the initial library. Comparison of the different variants to the original wild-type (WT) sequence for each library are showed in Figure 4. In all libraries, the negative control sequences consistently indicated a lack of binding, confirming the specificity of the antibodies for the antigen target. In three of the libraries, some variants successfully improved the affinity, by orders of magnitude in some cases (Table 4). For example, the binding of the peptide to the WT sequence in AAYL50 (Group 2) was not detectable (*>*200 *µ*M), but improved over 20-fold for variant Var4E (9.2 *±* 4.6 *µ*M). Two additional variants within this library, Var3D and Var3C, had similar affinities of 11.9 *±* 3.0 *µ*M and 9.4 *±* 4.2 *µ*M, respectively. Improvements were also observed in libraries one (AAYL49) and four (AAYL52), with variants reaching low and mid-10’s of micromolar affinity. In the remaining library (Group 3, AAYL51), the variants showed only marginal improvement in the binding affinities with only one variant, Var3F, with an affinity below 100 *µ*M. Overall, AbMAP had a hit rate of 82% on average in predicting sequences with a range of affinities toward the peptide antigen.

## Footnotes

1 The global alignment for sequence pairs was computed with no match or gap penalties

